# A molecular mechanism for high salt taste in *Drosophila*

**DOI:** 10.1101/2022.02.25.481885

**Authors:** Sasha A. T. McDowell, Molly Stanley, Michael D. Gordon

## Abstract

Dietary salt detection and consumption are crucial to maintaining fluid and ionic homeostasis. To optimize salt intake, animals employ salt-dependent activation of multiple taste pathways. Generally, sodium activates attractive taste cells, but attraction is overridden at high salt concentrations by cation non-selective activation of aversive taste cells. In flies, high salt avoidance is driven by both ‘bitter’ taste neurons and a class of glutamatergic ‘high salt’ neurons expressing pickpocket23 (ppk23). Although the cellular basis of salt taste has been described, many of the molecular mechanisms remain elusive. Here, we show that ionotropic receptor 7c (IR7c) is expressed in glutamatergic high salt neurons, where it functions with co-receptors IR76b and IR25a to detect high salt. Misexpression of IR7c in sweet neurons, which endogenously express IR76b and IR25a, confers responsiveness to non-sodium salts, indicating that IR7c is sufficient to convert a sodium-selective receptor to a cation non-selective receptor. Furthermore, the resultant transformation of taste neuron tuning switches potassium chloride from an aversive to an attractive tastant. This research provides insight into the molecular basis of monovalent and divalent salt taste coding and the full repertoire of IRs needed to form a functional salt receptor.

## INTRODUCTION

Salt is vital for various physiological processes such as electrolyte homeostasis, neuronal transmission, muscle contraction, and nutrient absorption. However, too much salt can produce ill effects. To balance need and excess, the valence of sodium for both mammals and insects differs based on salt concentration. Generally, sodium becomes increasingly attractive up to ∼100 mM and aversive beyond ∼250 mM^1–5^.

The appetitive-aversive dichotomy of salt is encoded by the balance of distinct salt-sensitive taste pathways: sodium-specific cells that drive consumption of low salt, and distinct cation non-selective cells that override attraction at higher concentrations to mediate aversion^1–9^. In mice, a dedicated population of taste receptor cells (TRCs) that expresses epithelial sodium channel (ENaC), specifically senses sodium, and mediates behavioral attraction to low NaCl^ref2^. On the other hand, high salt recruits the two main aversive taste pathways – bitter and sour – to promote avoidance^3^. Unlike each of the other primary taste modalities, molecular sensors for high salt remain unclear.

Innately attractive and aversive pathways have also been co-opted for low and high salt coding in *Drosophila*. Sweet gustatory receptor neurons (GRNs), labelled by gustatory receptor 64f (Gr64f), respond selectively to sodium and promote salt consumption^4^. A recently defined GRN population expressing IR94e also displays sodium-selective activation and may have a minor impact on attraction^4^. Conversely, high concentrations of any salt activate two aversive GRN populations: bitter GRNs labelled by Gr66a, and a population of ppk23-expressing glutamatergic GRNs (ppk23^glut^)^4,6,10^. Although ppk23 is a member of the ENaC family, *ppk23* is not required for high salt responses in ppk23^glut^ GRNs^4^. However, *IR25a* and *IR76b* are necessary for both the sodium-selective salt responses of sweet GRNs and the cation non-selective responses of ppk23^glut^ GRNs^4,5,11^.

The IR25a and IR76b co-receptors are broadly expressed in chemosensory tissues^12,13^. Although there are notable exceptions^14–18^, IR-dependent taste responses typically require both IR25a and IR76b. This includes roles in detecting salts^4^, carbonation^13^, fatty acids^19^, calcium^10^ and acids^20,21^. Given the evidence for heteromeric assembly of olfactory IR channels from two co-receptors and a more specific tuning IR subunit, it is likely that each distinct IR25a/IR76b-dependent taste function is mediated by these co-receptors complexing with an additional IR that confers specific tuning^13,19,22^. This notion was suggested by the requirement of IR62a in calcium detection in Ppk23 GRNs^10^. However, misexpression of IR62a in other IR25a/76b-containing GRNs failed to confer calcium sensitivity^10^. Thus, identifying the full composition of a functional IR taste receptor has remained elusive.

In this study, we show that IR7c acts with IR25a and IR76b to form a functional high salt receptor. IR7c is expressed in a subset of labellar ppk23^glut^ GRNs that is activated by both monovalent and divalent salts. However, *IR7c* mutants specifically lack physiological responses and behavioral aversion to high concentrations of monovalent salts. Moreover, ectopic expression of IR7c endows sweet neurons with the ability to sense non-sodium salts and overturns the flies’ innate KCl aversion in favor of attraction. These findings describe the first heteromultimeric IR complex that is both necessary and sufficient for salt sensing. They also provide insight into the elusive molecular nature of high salt detection.

## RESULTS

### IR7c functions in high salt taste neurons

We first identified IR7c as a candidate high salt IR based on its reported expression in taste neuron projections resembling those of ppk23^glut^ GRNs^4,13^. Using a *Gal4* knock-in to the *IR7c* locus (*IR7c*^*Gal4*^), we observed one IR7c-expressing GRN in most to all L-type sensilla and a few S-type and I-type sensilla on the labellum (Fig. 1A, B). Co-labelling of *IR7c*^*Gal4*^ with a *LexA* reporter for vesicular glutamate transporter (*VGlut-LexA*) revealed that all IR7c neurons are positive for VGlut (Fig. 1C). Since all glutamatergic GRNs on the labellum express ppk23, we also confirmed overlap between *IR7c*^*Gal4*^ and *ppk23-LexA* reporter expression (Fig. 1D). Therefore, IR7c is expressed within a subset of the ppk23^glut^ population. As expected, IR7c GRN axons project to the taste processing region of the brain called the subesophageal zone (SEZ) in a pattern resembling those of ppk23^glut^ (Fig. 1E). We also noted 1-2 IR7c-expressing GRNs per leg and observed projections to the ventral nerve cord consistent with those previously reported for ppk23^4,23^ (Fig. S1).

**Figure 1:**
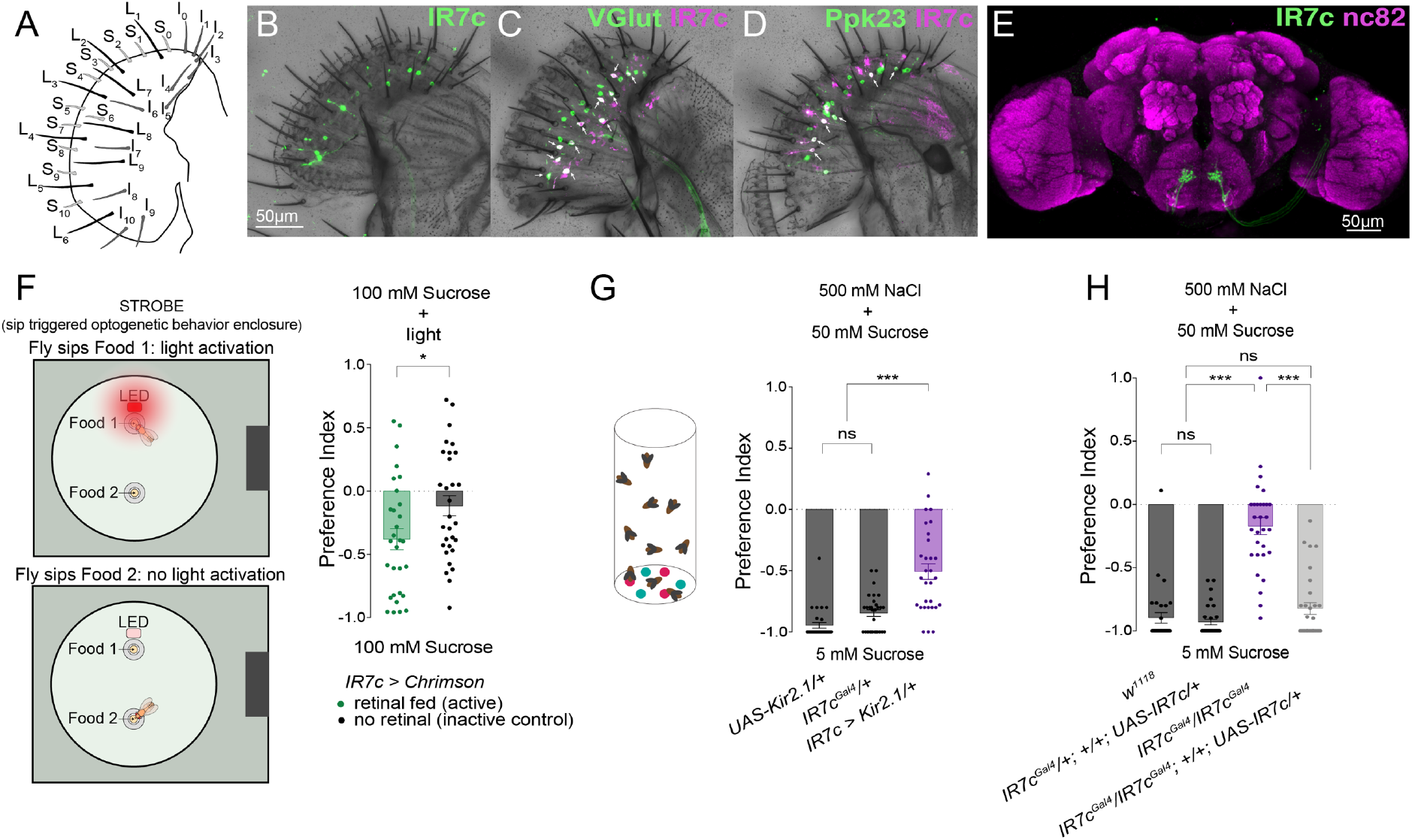
IR7c labels a high salt GRN population. **(A)** Schematic representation of the fly labellum. **(B-D)** Single labellar palps immunolabelled for *IR7c*^*Gal4*^ driving membrane tethered GFP (green) (B) or tdTomato (magenta) in combination with GFP (green) under the control of *vGlut-LexA* (C) or *ppk23-LexA* (D). Arrows indicate overlapping expression. **(E)** Immunofluorescence of IR7c GRN projections targeting the SEZ. **(F)** Schematic representation of the closed-loop sip triggered optogenetic behavior enclosure (STROBE; left) and preference indices for flies expressing CsChrimson in IR7c GRNs and fed retinal (green) or not fed retinal (grey). Positive values indicate preference for the light-triggering food. n = 30 flies per condition. **(G)** Preference indices for flies expressing Kir2.1 in IR7c GRNs (purple) and controls (grey) in the binary choice feeding assay. Positive values indicate preference for 500 mM NaCl plus 50mM sucrose; negative values indicate preference for 5 mM sucrose alone. n = 30 groups of ∼10 flies each. **(H)** Preference indices for *IR7c* mutants (purple), controls (dark grey) and rescue (light grey) in the high salt aversion binary choice assay. n = 30 groups of ∼10 flies each. Bars in all panels represent mean ± SEM. Asterisks denote significant difference between groups by unpaired two-tailed t-test (F) or one-way ANOVA with Tukey’s post hoc test (G, H), *p<0.05, ***p<0.001.

To determine whether IR7c neurons display functional properties equivalent to those of ppk23^glut^ GRNs, we first expressed CsChrimson under the control of *IR7c*^*Gal4*^ and tested the effect of closed-loop activation in the sip-triggered optogenetic behavior enclosure (STROBE)^24^ (Fig. 1F). Flies had access to feed on two identical sources of 100 mM sucrose, but interactions with one of the options triggered red LED illumination. Optogenetic activation of IR7c neurons prompted aversion of the light-triggering food, compared to control flies of the same genotype that were not fed the obligate CsChrimson cofactor all-*trans*-retinal (Fig. 1F). We next silenced IR7c GRNs using the Kir2.1 potassium channel and measured high salt avoidance using a dye-based binary feeding assay (Fig. 1G). Control flies avoided the high salt food option (500 mM NaCl + 50 mM sucrose) in favor of sucrose alone at a lower concentration, but flies with silenced IR7c GRNs showed significant impairment in their high salt aversion. Thus, IR7c GRNs carry negative valence and are required for normal avoidance of high salt.

Since IR76b and IR25a are both necessary for high salt detection by ppk23^glut^ GRNs^4^, we postulated that IR7c could be a more specific IR subunit that completes a functional high salt receptor. To detect a role for IR7c in high salt taste, we measured high salt avoidance in *IR7c*^*Gal4*^ mutants. As predicted, we found that *IR7c* mutants exhibit significant reduction in high salt avoidance, which was restored to control levels by the cell-type specific expression of *IR7c* (Fig. 1H).

### IR7c mediates a high salt response to NaCl and KCl

To more fully characterize IR7c’s role in salt detection and behavior, we expressed GCaMP7f under the control of *IR7c*^*Gal4*^ and imaged IR7c GRN axon terminals in the SEZ while stimulating the labellum for 5 seconds with increasing concentrations of NaCl and KCl (Fig. 2A, B). IR7c GRNs responded dose-dependently to increasing concentrations of both NaCl and KCl, exhibiting the lack of sodium selectivity characteristic of aversive high salt cells (Fig. 2C). Moreover, this response was lost in *IR7c* mutants and rescued by cell-type specific expression of *IR7c* (Fig. 2D, E), further confirming that IR7c labels high salt neurons and is essential for their responses to NaCl and KCl.

**Figure 2:**
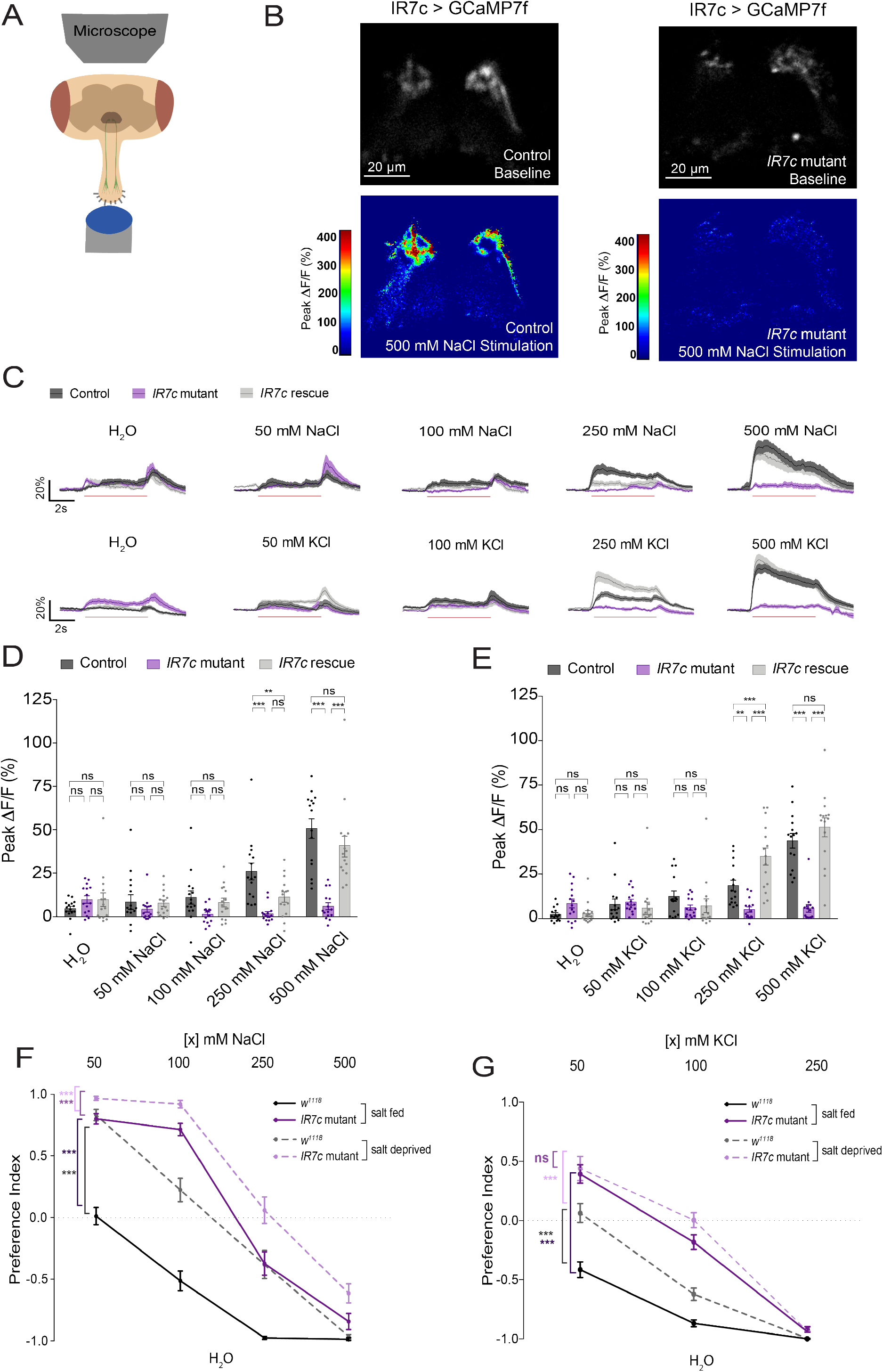
IR7c mediates behavioral avoidance of NaCl and KCl. **(A)** A schematic representation of the calcium imaging preparation. **(B)** Representative heatmaps showing IR7c GRNs stimulated with 500 mM NaCl in control (left) and *IR7c* mutant (right) flies. **(C)** Time traces of GCaMP7f signal in IR7c GRNs following stimulation with increasing concentrations of NaCl (top) and KCl (bottom). Trace lines and shaded regions represent mean ± SEM. Red line beneath traces indicates 5 s stimulation. Black trace lines denote the control genotype (*IR7c*^*Gal4*^/+ background), purple trace lines denote the *IR7c* mutants, and the light grey trace lines denote the *IR7c* rescue genotype. **(D, E)** Peak fluorescence changes during each stimulation. Bars represent mean ± SEM, n = 15 for each stimulus. Black dots (control), purple dots (*IR7c* mutants) and light grey dots (*IR7c* rescue) indicate values for individual replicates with n = 15 for each stimulus. Asterisks indicate significant difference between groups by two-way ANOVA with Tukey’s post hoc test, **p<0.01, ***p<0.001. (**F, G)** Preference indices for *IR7c* mutant (purple) and isogenic *w*^*1118*^ control flies (black) in a binary choice assay under salt fed (solid lines) or salt deprived (dashed lines) conditions. Positive values indicate preference for NaCl (E) or KCl (F) concentration on the x axis; negative values indicate preference for water. Dots represent mean ± SEM with n = 29-30 groups of ∼10 flies each for each genotype and condition. Asterisks denote significant difference between groups by two-way ANOVA with Bonferroni correction, ***p<0.001.

To understand how IR7c-mediated dose-dependent salt responses translate into behavior, we conducted NaCl and KCl binary-choice assays to assess flies’ preference for increasing concentrations of salt versus water. *w*^*1118*^ control flies that had been kept under salt fed conditions prior to the assay (three days with food containing 10 mM NaCl) to maximize salt avoidance, chose water and 50 mM NaCl equally. However, as salt concentration increased, flies showed dose-dependent salt avoidance (Fig. 2F). *IR7c* mutants kept under the same salt fed conditions displayed a dramatic reduction in salt aversion, although they still avoided the highest concentrations of NaCl, reflecting the existence of IR7c-independent salt avoidance mechanisms (Fig. 2F). Consistent with our past results, control flies that had been deprived of salt for three days showed significantly less salt aversion than salt fed flies of the same genotype (Fig. 2F). Nonetheless, salt deprived *IR7c* mutants exhibited a further reduction in avoidance compared to both salt deprived controls and salt fed *IR7c* mutants (Fig 2F). The results of the KCl binary-choice assay mirrored the NaCl behavioral findings, with the exception that there was no significant difference between salt fed and salt deprived *IR7c* mutants (Fig. 2G).

### IR7c is essential for IR7c GRN monovalent salt detection

Our next step was to determine the specific tuning of IR7c by conducting calcium imaging with a broader panel of stimuli encompassing different taste modalities (Fig. 3A) and different salt species (Fig. 3B). IR7c GRNs did not respond to sugar, bitter compounds, acetic acid, or a mixture of amino acids (AAs), demonstrating the specificity of these neurons for salt (Fig. 3C). IR7c GRN salt tuning, however, was broad, with strong responses to all salt species tested (Fig. 3D). Interestingly, while *IR7c* mutants showed complete loss of responses to all monovalent salts tested (NaCl, KCl, NaBr, CsCl), there was no effect on the detection of CaCl_2_ and MgCl_2_ (Fig. 3D). Since the anion species appeared to have no impact, we conclude that IR7c is specifically required for the detection of monovalent cations.

**Figure 3:**
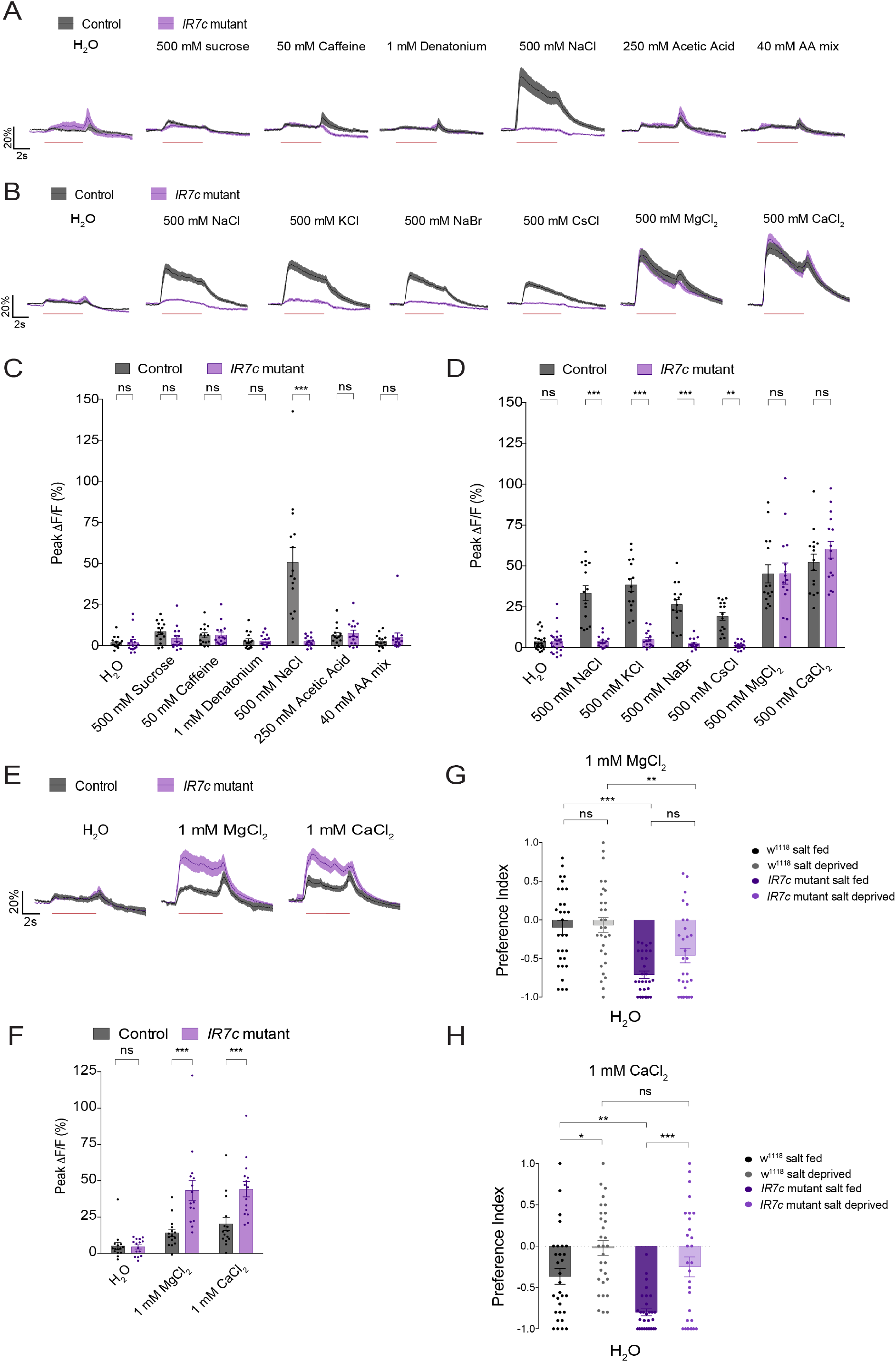
IR7c is essential for IR7c GRN monovalent salt detection. **(A-B)** Traces of GCaMP7f signal in IR7c GRNs of *IR7c* mutants (purple) and controls (grey), following stimulation with the indicated tastants. **(C, D)** Peak fluorescence changes during each stimulation. n = 15-23 for each stimulus. Black dots (control) and purple dots (*IR7c* mutants) indicate values for individual replicates. **(E)** Traces of GCaMP7f signal in IR7c GRNs following stimulation with 1 mM MgCl_2_ and 1 mM CaCl_2_. **(F)** Peak fluorescence changes during each stimulation, n = 15 for each stimulus. **(G, H)** Preference indices for *IR7c* mutants (purple) and controls (grey) in a binary choice assay, under salt fed (darker shades) and salt starved (lighter shades) conditions. Positive values indicate preference for the 1 mM MgCl_2_ (G) or 1 mM CaCl_2_ (H) negative values indicate preference for water. n = 30 groups of ∼10 flies each for each genotype and condition. All bars and traces represent mean ± SEM. Asterisks indicate significant difference between groups by two-way ANOVA with Sidak’s post hoc test (C, D, F) or one-way ANOVA with Tukey’s post hoc test (G, H), *p<0.05, **p<0.01, ***p<0.001.

Amongst the individual trials of 500 mM CaCl_2_ and MgCl_2_ stimulation, we noticed the salt responses in *IR7c* mutants to be somewhat higher than those of controls. However, this difference was difficult to tease apart due to a ceiling effect. Given that calcium is known to be detected at low concentrations^10^, we decided to examine the response of IR7c GRNs to low concentrations of divalent salts (Fig. 3E). IR7c GRNs showed subtle but consistent onset and removal responses to 500 mM CaCl_2_ and MgCl_2_, and 1 mM concentrations of both salts also elicited two peaks of calcium activity, albeit slightly less defined. *IR7c* mutants displayed a significantly enhanced onset peak, whilst the removal peak remained similar (Fig. 3E, F). Thus, loss of IR7c actually potentiates divalent salt sensitivity in IR7c GRNs.

The effect of IR7c loss on divalent salt feeding behavior prompted us to run CaCl_2_ and MgCl_2_ binary-choice assays. We found that *w*^*1118*^ flies generally had little preference for or against 1 mM MgCl_2_ versus water, whether flies were salt fed with NaCl or salt deprived (Fig. 3G). However, *IR7c* mutant flies had significantly stronger avoidance of 1 mM MgCl_2_ than controls under both conditions (Fig. 3G). Avoidance of CaCl_2_ by control flies was more strongly modulated by salt deprivation, with control flies exhibiting avoidance in the NaCl salt fed state but not when salt deprived (Fig. 3H). As with MgCl_2_, salt fed *IR7c* mutants displayed significantly stronger avoidance of 1 mM CaCl_2_; however, salt deprivation suppressed aversion to the point where it was not significantly different from controls (Fig. 3H).

One potential explanation for the potentiation of divalent salt responses in *IR7c* mutants is from increased availability of IR co-receptors to form a divalent salt receptor with a distinct tuning IR. Since IR62a has been implicated in calcium sensing^10^, we explored its involvement in divalent salt detection by IR7c GRNs. Calcium imaging of IR7c GRN divalent salt responses in *IR62a* mutant flies failed to reveal significant differences in the response of *IR62a* mutant flies to low or high concentrations of CaCl_2_ or MgCl_2,_ compared to controls (Fig. S2A, B). There was a trend towards reduced sensitivity to low divalent salt concentrations, including 50 mM CaCl_2_, which was previously shown to evoke IR62a-dependent electrophysiological responses in labellar taste sensilla, but the effect did not reach significance^10^ (Fig. S2C, D). We reasoned that perhaps IR62a and IR7c could be partially redundant in their functions. However, flies that were mutant for both *IR7c* and *IR62a* retained their low and high responses to the divalent salts tested (Fig. S2E, F). Thus, we conclude that divalent salt sensitivity in IR7c neurons can be mediated independently of IR7c and IR62a.

### All ppk23 GRNs require IR7c for monovalent salt detection

Given the likelihood of multiple high salt receptor complexes, we wanted to clarify the extent to which IR7c is responsible for salt responses across different GRN types in the labellum. We therefore used *ppk23-LexA, Gr66a-LexA* (bitter), and *Gr64f-LexA* (sweet) to perform calcium imaging of the main salt-sensitive GRN populations, and compared responses in *IR7c* mutants and isogenic controls (Fig. 4). Ppk23 GRNs displayed the same properties as the IR7c population, with *IR7c* essential for monovalent but not divalent high salt responses (Fig 4A, D, G). By contrast, bitter GRNs showed little activity to divalent salts and their monovalent salt responses were largely *IR7c*-independent, although a significant reduction was evident in response to 500 mM KCl (Fig 4B, E, H). This conclusion is also supported by our prior observation that bitter GRN salt responses are largely IR25a- and IR76b-independent^4^, and by co-labelling of *IR7c*^*Gal4*^ with *Gr66a-LexA*, which revealed little to no detectable overlap (Fig. S3A, B). Lastly and as expected, *IR7c* mutations had no effect on salt responses in sweet (Gr64f) GRNs (Fig. 4C, F, I). Together, these data suggest that IR7c is an obligate component of labellar IR receptor complexes mediating monovalent high salt detection.

**Figure 4:**
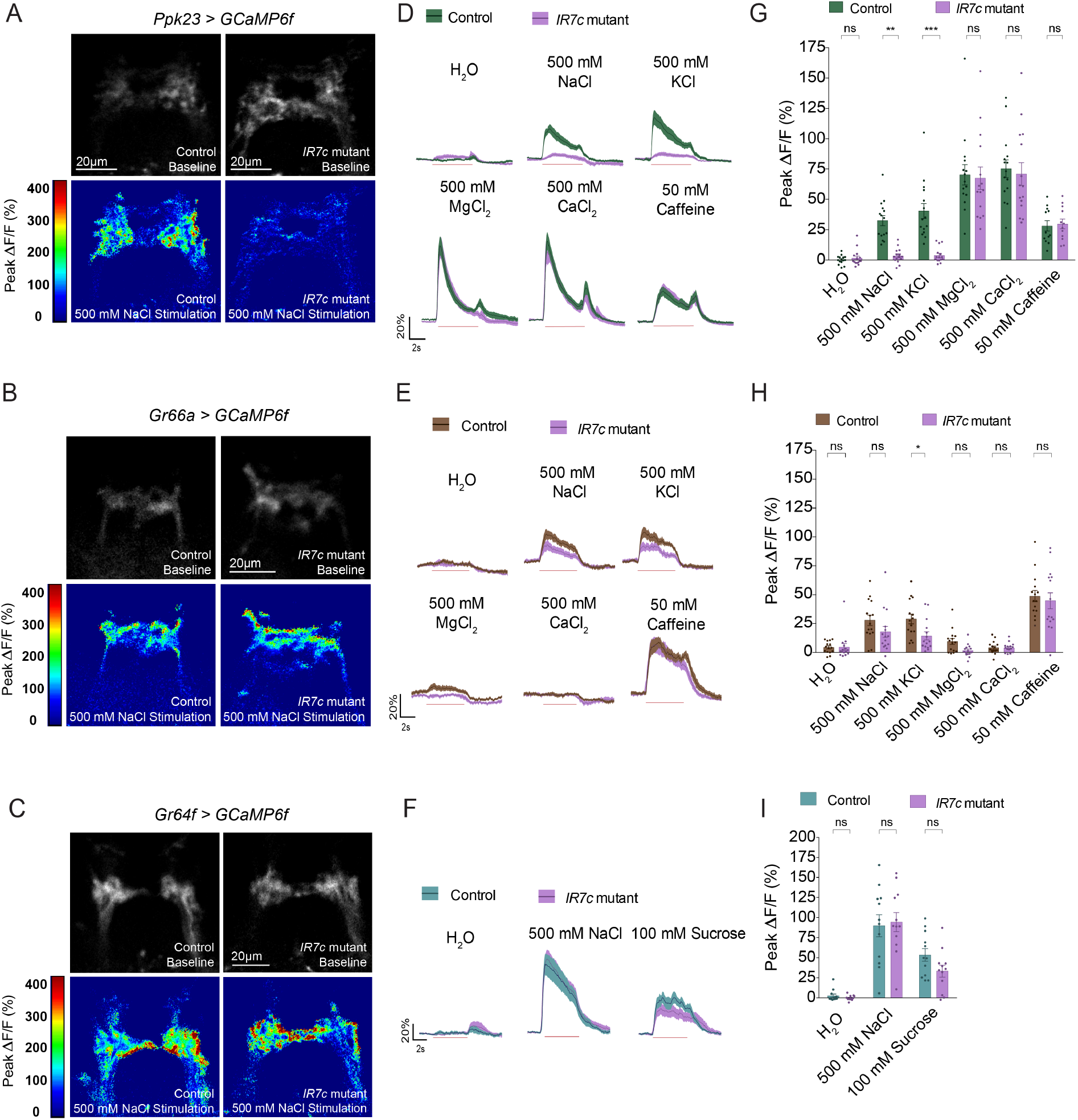
IR7c is essential for monovalent salt responses in Ppk23 GRNs. **(A-C)** Representative heatmaps of 500 mM NaCl-evoked activity for each indicated GRN type from isogenic control (left) and *IR7c* mutant (right) flies. **(D-F)** Traces of GCaMP6f signal for each indicated GRN type, following stimulation with the indicated tastants. Trace lines and shaded regions represent mean ± SEM. Red line beneath traces indicates a 5 s long stimulation. Green, brown and blue trace lines denote control genotypes (*IR7c*^*Gal4*^/+ background) and purple trace lines denote *IR7c* mutant genotypes. **(G-I)** Peak fluorescence changes during each stimulation. Bars represent mean ± SEM, n = 12-15 for each stimulus. Green, brown and blue dots (control), and purple dots (*IR7c* mutants) indicate values for individual replicates. Asterisks indicate significant difference between control and mutant responses for each stimulus by two-way ANOVA with Sidak’s post hoc test, *p<0.05, **p<0.01, ***p<0.001.

### IR7c forms a functional salt receptor with IR25a and IR76b

We next sought to define the full composition of the monovalent high salt receptor complex. First, we confirmed that IR76b and IR25a are required for salt responses in IR7c GRNs. Indeed, *IR76b* and *IR25a* mutants display a complete loss of salt evoked activity in IR7c neurons (Fig. 5A-D). In both cases, responses were rescued by cell-type specific rescue using *IR7c*^*Gal4*^. Therefore, IR76b and IR25a are necessary for both monovalent and divalent salt detection in IR7c GRNs, while IR7c is specifically required for monovalent salts.

**Figure 5:**
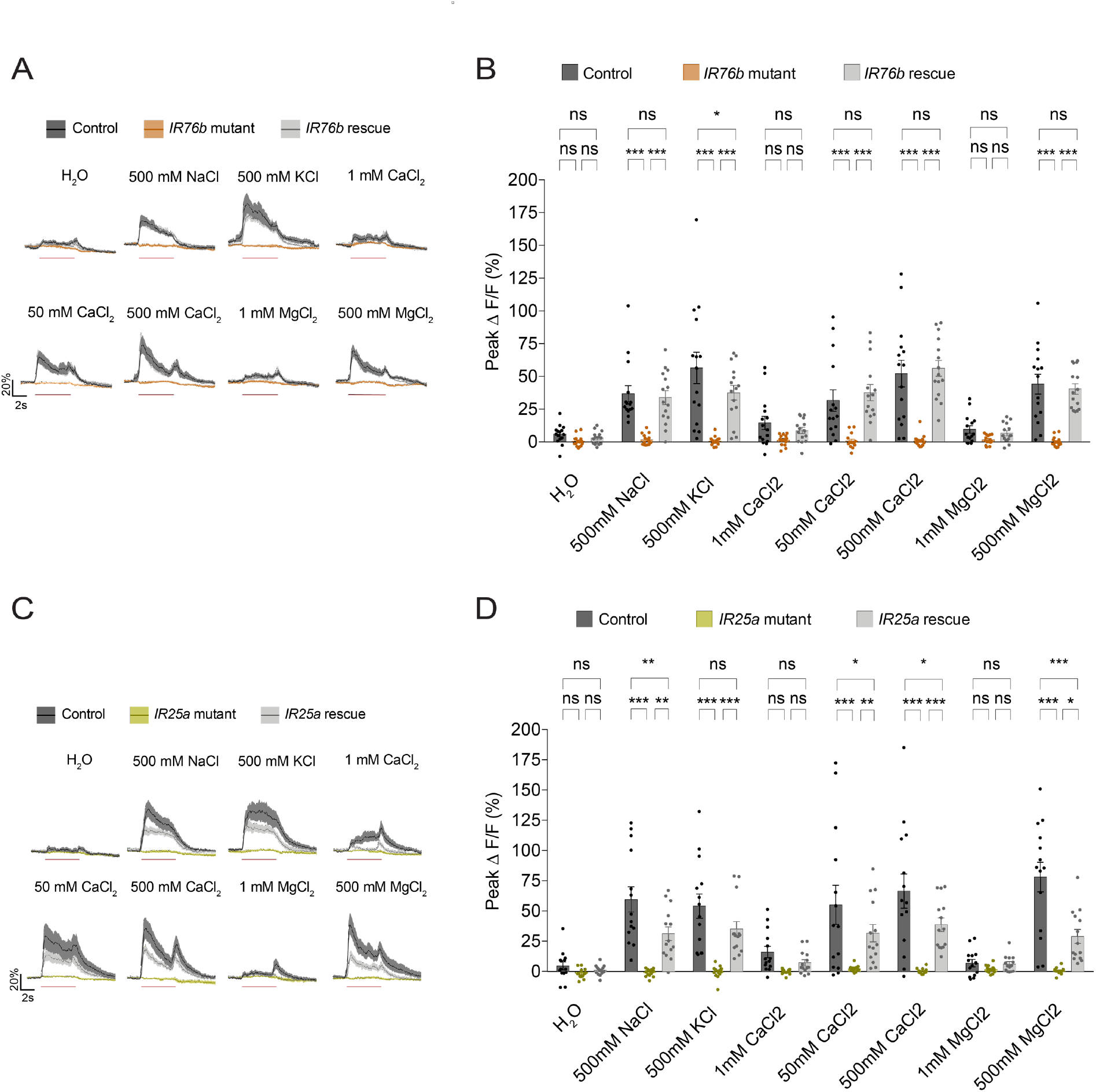
IR76b and IR25a are necessary for IR7c GRN salt responses. **(A)** Traces of GCaMP7f signal in IR7c GRNs of *IR76b* mutants (orange), heterozygous controls (black) and *IR76b* rescue flies (grey), following stimulation with the indicated tastants. **(B)** Peak fluorescence changes during each stimulation. Black dots (control), orange dots (*IR76b* mutant) and grey dots (*IR76b* rescue) indicate values for individual replicates. n = 15 for each stimulus. **(C)** Traces of GCaMP7f signal in IR7c GRNs of *IR25a* mutants (green), heterozygous controls (black) and *IR25a* rescue flies (grey), following stimulation with the indicated tastants. **(D)** Peak fluorescence changes during each stimulation. Black dots (control), green dots (*IR25a* mutant) and grey dots (*IR25a* rescue) indicate values for individual replicates. n = 13-14 for each stimulus. All bars and traces represent mean ± SEM. Asterisks indicate significant difference between control, mutant and rescue responses for each stimulus by two-way ANOVA with Sidak’s post hoc test, *p<0.05, **p<0.01, ***p<0.001.

To establish that IR25a, IR76b, and IR7c complex to form a functional high salt receptor, we aimed to reconstitute this receptor in a heterologous GRN type. Thus, we used *Gr64f-Gal4* to express IR7c in sweet neurons, which already express IR25a and IR76b^4,13,18,19,21^ (Fig. 6A). Strikingly, while sweet neurons normally show responses only to sodium salts, IR7c expression conferred sensitivity to high concentrations of CaCl_2_ and KCl (Fig. 6B, C). Thus, expression of IR7c was sufficient to convert sweet neurons from a sodium-specific ‘low salt’ cell to a cation non-selective ‘high salt’ cell type. Interestingly, this experiment also indicated that an IR7c-containing salt receptor is sufficient, but not necessary, for detection of high concentrations of divalent salts. Along with the observation that IR7c expression did not evoke low CaCl_2_ responses, this suggests that IR7c/25a/76b form a cation non-specific high salt receptor, but that more sensitive IR25a/76b-dependent divalent salt receptor(s) exist in IR7c GRNs. To further support IR7c’s sufficiency in the formation of a high salt receptor, we misexpressed IR7c in a second heterologous GRN population using *IR94e-Gal4*. Wild-type IR94e GRNs display minimal low sodium activation^4^, but we conducted the experiment in an *IR94e* mutant background, which shows no significant high salt-evoked activity (Fig S4A-C). Nonetheless, upon ectopic expression of *IR7c*, strong cation non-selective high salt responses emerged (Fig. S4A-C). Notably, reconstituting the IR7c high salt receptor in sweet or IR94e GRNs produced onset and offset peaks to calcium, suggesting that these dynamics are a property of the receptor and not specific to the IR7c GRN type.

**Figure 6:**
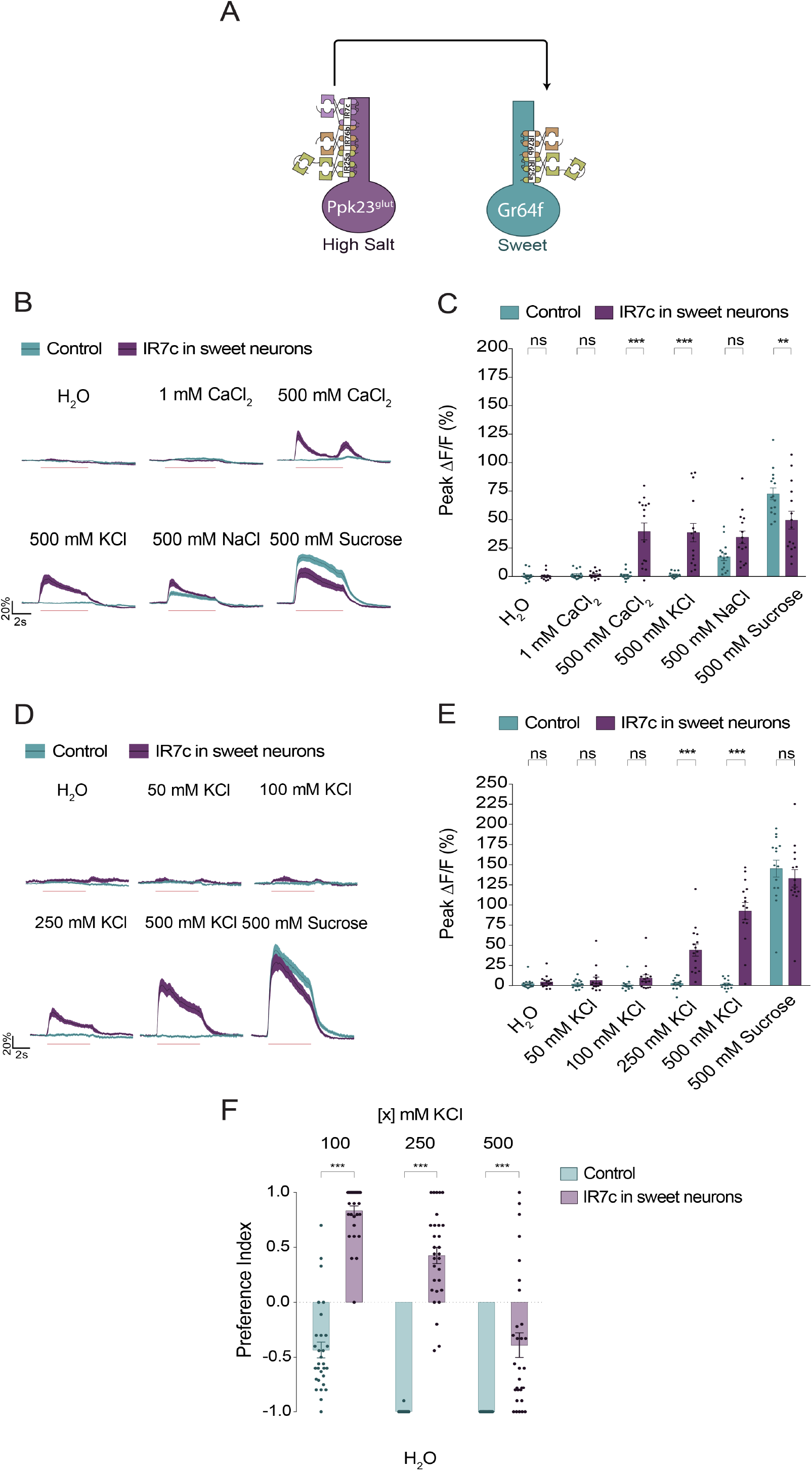
Ectopic IR7c expression in sugar sensing GRNs reconstitutes a high salt receptor. **(A)** A schematic representation of inducing a high salt taste receptor in sweet-sensing neurons. **(B, D)** Traces of GCaMP6f signal in Gr64f GRNs of flies with IR7c expression under the control of *Gr64f-Gal4* (purple) and isogenic controls (blue), following with the indicated tastants. **(C, E)** Peak fluorescence changes during each stimulation. Blue dots (control) and purple dots (flies with IR7c expression under the control of *Gr64f-Gal4*) indicate values for individual replicates. n = 15 for each stimulus. **(F)** Preference indices for flies with IR7c expression under the control of *Gr64f-Gal4* (purple) and an isogenic control (blue) in a binary choice assay. Positive values indicate preference for indicated KCl concentrations; negative values indicate preference for water. n = 30 groups of ∼10 flies each for each genotype. All bars and traces represent mean ± SEM. Asterisks indicate a significant difference between the two genotypes tested by two-way ANOVA with Sidak’s post hoc test, **p<0.01, ***p<0.001.

Further calcium imaging of sweet GRNs with IR7c overexpression revealed dose-dependent responses to KCl (Fig. 6D, E). We predicted that transforming the stimulus specificity of sweet GRNs, which evokes appetitive behavior, would impact flies’ feeding on non-sodium salts. KCl, like other non-sodium salts, is generally aversive at all concentrations (Fig. 6F). However, flies overexpressing IR7c in sweet GRNs were attracted to KCl at 100 mM and 250 mM and displayed significantly less aversion to 500 mM KCl (Fig. 6F). Therefore, broadening the salt tuning of sweet GRNs is sufficient to induce KCl attraction.

### IR62a antagonizes IR7c GRN salt responses

Having established that IR7c can confer high salt responses in Gr64f GRNs, we investigated whether IR62a may participate in detection of low CaCl_2_. Consistent with a previous report^10^, the sole overexpression of IR62a in sweet sensing neurons is insufficient to evoke calcium responses (Fig. 7A, B). When IR7c and IR62a were co-expressed in Gr64f GRNs, no new calcium responses were observed beyond what was seen for IR7c expression alone (Fig. 7C-D). Oddly, we noticed that including IR62a dampened IR7c-mediated KCl responses and completely abolished the response to CaCl_2_ (Fig. 6C). To understand the interaction between these two receptors further, we overexpressed IR62a in IR7c GRNs. Interestingly, this caused a drastic reduction in all IR7c GRN salt responses (Fig. 7E, F). Therefore, IR62a antagonizes IR7c’s function in salt detection, possibly through displacement of IR7c from the IR7c/25a/76b complex. Along with our observation that IR7c mutants display elevated calcium and magnesium responses, this suggests that IR-dependent tuning is highly sensitive to subunit dose.

**Figure 7:**
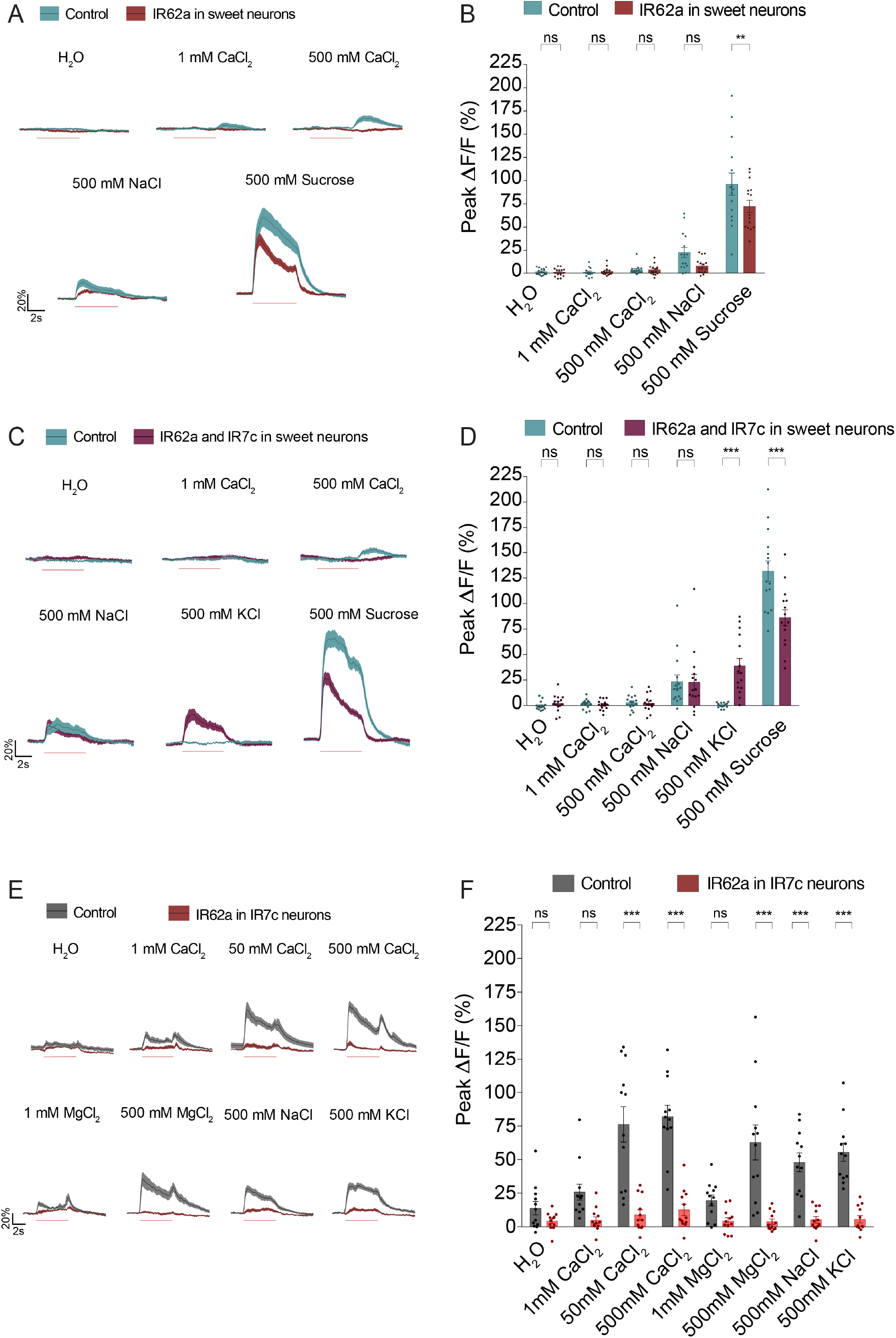
IR62a antagonizes IR7c GRN salt responses. **(A)** Traces of GCaMP6f signal in Gr64f GRNs of flies with IR62a expression under the control of *Gr64f-Gal4* (red) and isogenic controls (blue), following stimulation with the indicated tastants. **(B)** Peak fluorescence changes during each stimulation. Blue dots (control), and red dots (flies with IR62a expression under the control of *Gr64f-Gal4*) indicate values for individual replicates with an n = 15 for each stimulus. **(C)** Traces of GCaMP6f signal in Gr64f GRNs of flies with both IR7c and IR62a expression under the control of *Gr64f-Gal4* (purple) and isogenic controls (blue), following stimulation with the indicated tastants. **(D)** Peak fluorescence changes during each stimulation. Blue dots (control), and purple dots (flies with both IR7c and IR62a expression under the control of *Gr64f-Gal4*) indicate values for individual replicates with an n = 15 for each stimulus. **(E)** Traces of GCaMP7f signal in IR7c GRNs of flies with IR62a expression under the control of *IR7c*^*Gal4*^ (red) and isogenic controls (grey), following 5 s stimulation (red line) with the indicated tastants. **(F)** Peak fluorescence changes during each stimulation. Black dots (control), and red dots (flies with IR62a expression under the control of *IR7c*^*Gal4*^) indicate values for individual replicates. n = 12 for each stimulus. All bars and traces represent mean ± SEM. Asterisks indicate significant difference between responses of the different genotypes for each stimulus by two-way ANOVA with Sidak’s post hoc test, **p<0.01, ***p<0.001.

## DISCUSSION

The prior observation that IR76b and IR25a are necessary components of both sodium-specific and cation non-selective salt taste receptors raised the question of how differences in tuning are achieved in attractive and aversive salt-sensitive taste cells^4,5,11^. This study identifies IR7c as a critical component of the receptor for monovalent high salt taste, and provides insights into how monovalent and divalent salt avoidance is encoded in flies (Fig.8A)

**Figure 8:**
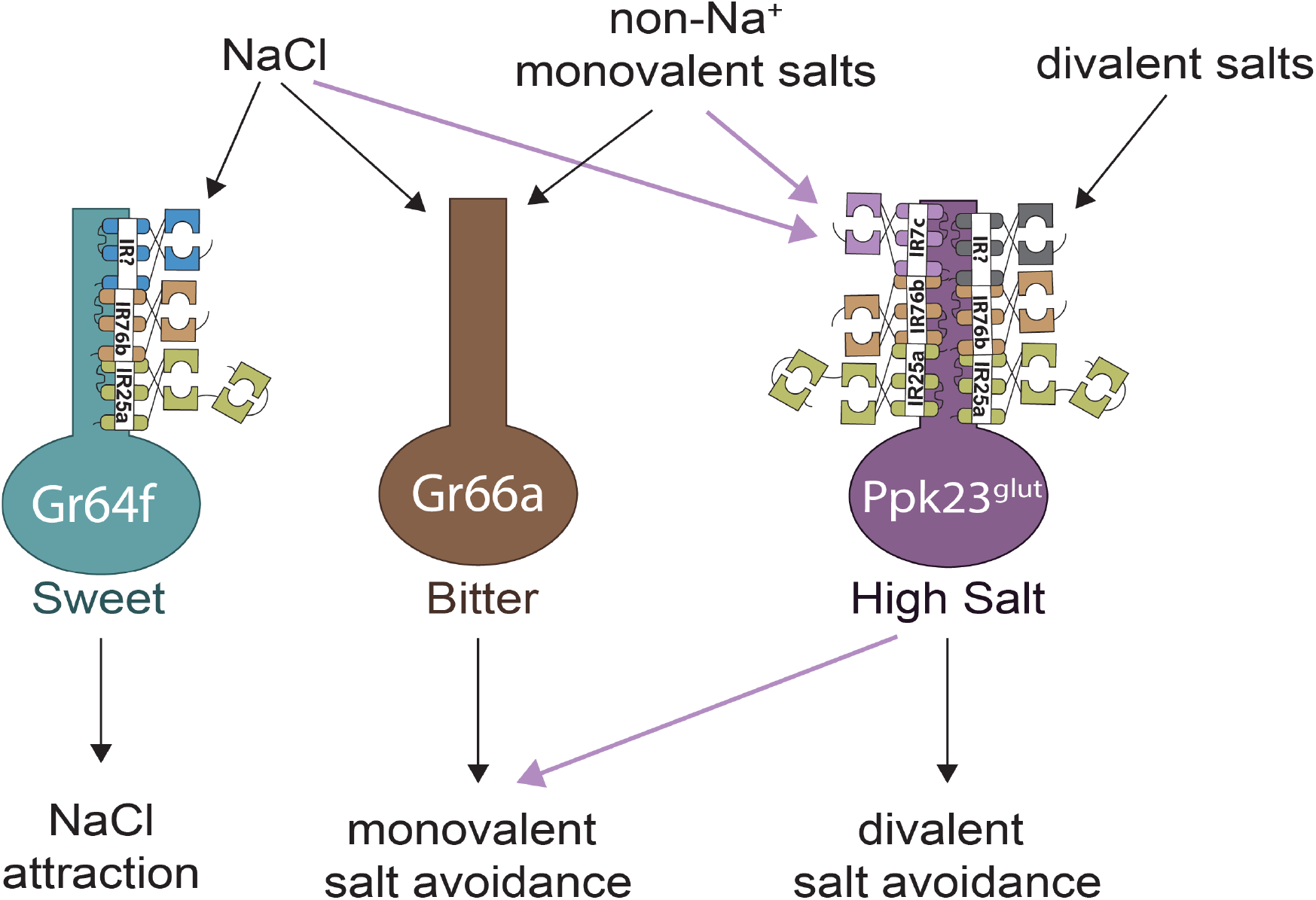
Molecular mechanisms for salt encoding across the main salt-sensitive GRN populations. Arrows indicate excitation that leads to subsequent behavioral consequences, with the roles for which IR7c is necessary for highlighted by purple arrows.

### Monovalent salt coding

Unlike the peripheral coding of bitter and sweet compounds, which follows a labeled-line organization^6,8,9,25^, monovalent salts are encoded by a combination of most to all GRN classes^4^. Although neural silencing of distinct GRN populations has previously revealed contribution of both bitter and ppk23^glut^ GRNs to high salt avoidance, *IR7c* mutants have afforded an additional tool to further probe the complexity of monovalent salt taste coding. In particular, the drastic change in behavior towards NaCl seen in *IR7c* mutants highlights the importance of glutamatergic GRNs in high salt avoidance, although IR7c-independent salt avoidance mechanisms clearly exist (Fig. 2F). These behavioral shifts are especially notable given that they are seen in the absence of the sugar typically used in high salt assays to incentivize consumption of the salt-containing option. Interestingly, *IR7c* mutants also display attraction towards 50 mM KCl (Fig. 2G), despite the lack of KCl-evoked activity in sweet GRNs (Fig. 6E). The source of this attraction is unclear, but could be derived from very weak activation of sweet neurons or osmotic effects on ppk28-expressing ‘water’ GRNs^26,27^.

We previously suggested that aversion via the ppk23^glut^ salt-specific pathway is suppressed by salt deprivation, thereby fine-tuning salt intake based on need^4^. Consistent with this model, we find that the avoidance of KCl is modulated by salt deprivation in control flies, but not in *IR7c* mutants (Fig. 2G). Thus, the IR7c salt taste pathway is necessary for modulation of KCl avoidance by salt need. However, modulation of NaCl avoidance by salt deprivation is only partially suppressed in *IR7c* mutants, as salt deprived mutants show marginally higher NaCl preference than those that have been previously salt fed (Fig. 2F). Together, these results suggest that sodium-specific attractive salt taste is also modulated (in this case enhanced) by salt deprivation. Finally, it is notable that even among salt-deprived flies, *IR7c* mutants show significantly reduced salt avoidance compared to controls. This suggests that our 3-day salt deprivation did not fully suppress the IR7c-dependent pathway to the equivalence of IR7c loss. Whether longer deprivation could more fully suppress this pathway is yet to be tested.

### Divalent salt coding

IR7c is not necessary for divalent responses in IR7c GRNs (Fig. 3D, 3F) or the broader ppk23 population (Fig. 4G); however, it is sufficient to confer high CaCl_2_ responses in Gr64f neurons (Fig 6C). This suggests that IR7c is a cation non-specific high salt receptor, but that more specific divalent salt receptor(s) exist in IR7c GRNs and possibly the broader ppk23 population. Since consumption of small concentrations of divalent salts can be detrimental to a fly’s fitness^10^, receptors that are sensitive to low divalent concentrations are needed to mediate avoidance. This could explain the functional redundancy of divalent receptors within IR7c GRNs. Surprisingly, loss of IR7c causes increased divalent taste responses and aversion (Fig. 3E-H). Perhaps, since IR76b and IR25a are necessary for these divalent responses (Fig. 5), loss of IR7c “frees up” these coreceptors to form more divalent receptors. Alternatively, *IR7c* mutant flies may consume more salt, effectively creating a “salt fed” condition in which IR7c GRNs may be modulated to increase their aversion towards all salts including divalent ones. However, prior calcium imaging of ppk23^glut^ GRNs did not reveal modulation at the level of GRN activity^4^.

Based on our findings, IR62a is not responsible for calcium responses in IR7c GRNs. Interestingly, the potentiation of low divalent salt responses observed in IR7c mutants disappeared in the IR62a and IR7c double mutant, indicating that IR62a may play a role in this effect. However, overall, our IR62a results contrast with a previous report that 50 mM CaCl_2_ evoked IR62a-dependent electrophysiological activity in S-type sensilla^10^. IR7c GRNs innervate all L-type sensilla and only a few S-type sensilla (Fig. 1B). Therefore, it is possible that IR62a functions in calcium detection within IR7c-negative ppk23 GRNs. Silencing IR7c GRNs and imaging the ppk23 population could address this possibility. Why flies would have different divalent cation receptors within the IR7c subpopulation of ppk23 GRNs versus outside of it is thought-provoking. One clue may exist in the discovery that IR62a antagonizes IR7c GRN salt responses (Fig. 7E, F). How this occurs is unclear, but, plausibly, competition for IR76b and IR25a coreceptors could explain the phenomenon. Regardless, this finding makes it unlikely that IR7c and IR62a receptors function within the same neurons, unless their expression levels are tightly controlled to ensure appropriate stoichiometry among different receptors. Moreover, perhaps having divalent receptor(s) outside the IR7c subpopulation of ppk23 GRNs could allow for discrimination between different salt species.

One physiological curiosity relating to divalent salt taste is the two calcium activity peaks corresponding to onset and removal. Similar kinetics have been observed in bitter GRN responses to bitter substances^28,29^ and in sweet GRN acid responses^18^. As in bitter taste^28,29^, we find that the same receptor, in our case IR7c, can mediate both onset and removal peaks. It will be interesting to examine the contribution to taste coding afforded by these temporal dynamics, and whether both peaks carry the same valence, as suggested for attractive acids^18^, or different valence, as suggested for bitter compounds^28^.

### Minimum IRs required to form a taste receptor

Several studies have described specific sets of three IRs that are necessary for diverse roles in chemosensation^10,13,19,22^ and hygrosensation^30–33^. In addition to demonstrating necessity of IR7c, IR76b, and IR25a in monovalent high salt detection, we showed that misexpressing IR7c in two different neuronal populations already expressing IR76b and IR25a leads to reconstitution of a high salt receptor (Fig. 6, S4). A similar experiment was previously done with IR7a, which confers acetic acid sensitivity to L-type sensilla when misexpressed in Gr5a GRNs^16^. However, although IR7a appears to function independent of IR25a and IR76b, IR7a was insufficient to confer acetic-acid-evoked conductance to *Drosophila* S2 cells, suggesting that perhaps additional co-receptors are required^16^. Our experiments suggest both necessity and sufficiency of an IR7c/25a/76b complex in mediating monovalent high salt taste. It is formally possible that, by chance, other IR subunits necessary for this high salt receptor are expressed in both IR94e and Gr64f GRNs. However, we consider this possibility improbable. Instead, we postulate that IR25a, IR76b and IR7c constitute a minimum set of IRs to form a salt taste receptor.

Our results also beg the question of whether there is an uncharacterized tuning IR that is expressed in sweet GRNs and cooperates with IR25a and IR76b to form a sodium-selective salt receptor. If IR25a and IR76b were the only receptors required for NaCl sensing, then the loss of IR7c would leave these coreceptors intact and capable of generating NaCl responses. Since this was not observed (Fig. 2D), it is likely that sodium-specific taste requires an IR not expressed in IR7c GRNs. Moreover, although sweet GRNs respond in a dose-dependent manner towards NaCl, they appear to have a lower threshold for NaCl detection than IR7c GRNs^4^. Understanding the biophysical mechanisms by which different subunits confer changes in both ion selectivity and sensitivity to a salt taste receptor will be illuminating.

To date, a high salt taste sensor has not been unequivocally identified in mammals. Since the primary effectors of T2R signaling, PLC-β2 and TRPM5, are necessary for high-salt sensing in bitter TRCs, it is likely one or more of the approximate 30 T2Rs are sensitive to high salt concentrations^3^. There is also evidence that carbonic anhydrase 4, an enzyme involved in buffering the pH around TRCs, may function to translate external salt into local pH changes in turn activating the intrinsic sour-TRC mechanism^3^. Although IRs are not conserved in mammals^34^, uncovering their general principles as salt sensors may provide insight into mammalian high salt detection mechanisms.

## ACKNOWLEDGEMENTS

We thank Kiereth Atariwala for assistance with binary choice feeding assays and members of the Gordon lab for comments on the manuscript. We also thank the Bloomington Stock Centre for fly stocks. This work was funded by the Canadian Institute of Health Research (CIHR) operating grant FDN-14842. M.D.G. is a Michael Smith Foundation for Health Research Scholar

## AUTHOR CONTRIBUTIONS

S. A. T. M and M. D. G. conceived the project and wrote the manuscript. S. A. T. M performed all experiments. M. S. performed pilot GCaMP imaging and gave experimental advice. M. D. G. supervised the project.

## DECLARATION OF INTERESTS

The authors declare no competing interests

## SUPPLEMENTAL INFORMATION

Supplemental information includes four figures

## MATERIALS AND METHODS

### Flies

Flies were raised on standard cornmeal diet at 25°C in 70% humidity. All experimental flies were 2-10 days mated females unless otherwise stated. Information on flies generated in this study and the genotypes for each experiment are given below. Information on other flies used in this study can be found in the Key Resources Table.

### Generation of IR7c and IR94e lines

The *IR7c*^*Gal4*^ line was created by deleting most of the coding DNA sequence (CDS) and replacing it with Gal4::VP16 using CRISPR/Cas9-mediated genome editing with homology-dependent repair. Upstream gRNA sequence GCAACATCGTGTTTCATCGG[TGG] and downstream gRNA sequence GATTTGTGGTCAAGATCTCC[AGG] were cloned into the U6 promoter plasmid. Upstream and downstream homology arms of *IR7c* were amplified by PCR and subcloned into the pUC57-Kan vector with a *Gal4::VP16-RFP* cassette containing Gal4::VP16, 3xP3-RFP, and two loxP sites. IR7c-targeting gRNAs and hs-Cas9 were microinjected with the donor plasmid into a *w*^*1118*^ control strain. F1 flies carrying the selection marker were validated by genomic PCR and sequencing. The resulting deletion was 1919bp, from -12^th^ nucleotide relative to ATG to -65^th^ nt relative to stop codon of *IR7c*. The entire procedure was performed by WellGenetics Inc. (Taipei City, Taiwan) using a modified version of previously published methods^35^.

The *IR94e*^*LexA*^ line was created using a similar method as *IR7c*^*Gal4*^ with the exception that the IR94e CDS was deleted and replaced by *nls-LexA::P65*. The gRNA sequences were TGCCCAAAGTGGATCCTGAG[CGG] and TTCCAGCAGCCAAACTAGCG[AGG] The upstream and downstream homology arms of *IR94e* were amplified by PCR and subcloned into pUC57-Kan vector with *nls-LexA::P65-RFP* cassette containing *nls-LexA::P65-RFP*, two loxP sites, and 3xP3-RFP. The resulting deletion was 1843bp, from the start codon “ATG” to -49^th^ nucleotide relative to the *Ir94e* stop codon. The entire procedure was performed by WellGenetics Inc. (Taipei City, Taiwan) using a modified version of previously published methods^35^.

The *UAS-IR7c* transgenic line was created by synthesizing the coding sequence of *IR7c* and subcloning into the NotI site of the PUAST-attB vector. Synthesis and cloning were performed by Bio Basic Inc. (Ontario, Canada). The transformation vector was injected into *w*^*1118*^ embryos for PhiC31c-mediated integration into the attP2 site. Injections were performed by Rainbow Transgenic Flies Inc. (California, USA).

### Fly genotypes by figure

Figure 1:

- *IR7c*^*Gal4*^*/IR7c*^*Gal4*^; *+/+; UAS-CD8::GFP/+*
- *IR7c*^*Gal4*^*/+; vGlut-LexA/UAS-tdTomato; LexAop2-mCD8:GFP/+*
- *IR7c*^*Gal4*^*/IR7c*^*Gal4*^; *ppk23-LexA/UAS-CD8::tdTomato; LexAop2-mCD8:GFP/+*
- *IR7c*^*Gal4*^*/IR7c*^*Gal4*^; *+/+; UAS-CD8::GFP/+*
- *IR7c*^*Gal4*^*/IR7c*^*Gal4*^; *UAS-CsChrimson/+; +/+*
- *+/+; +/+; UAS-Kir2*.*1/+*
- *IR7c*^*Gal4*^*/+; +/+; +/+*
- *IR7c*^*Gal4*^*/+; +/+; UAS-Kir2*.*1/+*
- *w*^*1118*^
- *IR7c*^*Gal4*^*/+; +/+; UAS-IR7c/+*
- *IR7c*^*Gal4*^*/IR7c*^*Gal4*^; *+/+; +/+*
- *IR7c*^*Gal4*^*/IR7c*^*Gal4*^; *+/+; UAS-IR7c/+*

Figure 2:

- *IR7c*^*Gal4*^*/+; +/+; UAS-GCaMP7f/UAS-GCaMP7f*
- *IR7c*^*Gal4*^*/IR7c*^*Gal4*^; *+/+; UAS-GCaMP7f/+*
- *IR7c*^*Gal4*^*/IR7c*^*Gal4*^; *+/+; UAS-GCaMP7f/UAS-IR7c*
- *IR7c*^*Gal4*^*/IR7c*^*Gal4*^; *+/+; +/+*
- *w*^*1118*^

Figure 3:

- *IR7c*^*Gal4*^*/+; +/+; UAS-GCaMP7f/UAS-GCaMP7f*
- *IR7c*^*Gal4*^*/IR7c*^*Gal4*^; *+/+; UAS-GCaMP7f/+*
- *IR7c*^*Gal4*^*/IR7c*^*Gal4*^; *+/+; +/+*
- *w*^*1118*^

Figure 4:

- *IR7c*^*Gal4*^*/IR7c*^*Gal4*^; *ppk23-LexA/ LexAOp-GCaMP6f; +/+*
- *IR7c*^*Gal4*^*/IR7c*^*Gal4*^; *Gr66a-lexA/ LexAOp-GCaMP6f; +/+*
- *IR7c*^*Gal4*^*/IR7c*^*Gal4*^; *+/LexAOp-GCaMP6f; Gr64f-LexA/+*

Figure 5:

- *IR7c*^*Gal4*^*/+; UAS-GCaMP7f/+; IR76b*^*1*^
- *IR7c*^*Gal4*^*/+; UAS-GCaMP7f/+; IR76b*^*1*^*/IR76b*^*2*^
- *IR7c*^*Gal4*^*/+; UAS-GCaMP7f/UAS-IR76b; IR76b*^*1*^*/IR76b*^*2*^
- *IR7c*^*Gal4*^*/+; IR25a*^*1*^*/+; UAS-GCaMP7f/+*
- *IR7c*^*Gal4*^*/+; IR25a*^*1*^*/IR25a*^*2*^; *UAS-GCaMP7f/+*
- *IR7c*^*Gal4*^*/+; IR25a*^*1*^*/IR25a*^*2*^, *UAS-IR25a; UAS-GCaMP7f/+*

Figure 6:

- *+/+; Gr64f-Gal4, UAS-GCaMP6f/+; +/+*
- *+/+; Gr64f-Gal4, UAS-GCaMP6f/+; UAS-IR7c/+*
- *+/+; Gr64f-Gal4/+; +/+*
- *+/+; Gr64f-Gal4/+; UAS-IR7c/+*

Figure 7:

- *+/+; Gr64f-Gal4, UAS-GCaMP6f/+; +/+*
- *+/+; Gr64fGal4, UAS-GCaMP6f/UAS-IR62a; +/+*
- *+/+; Gr64f-Gal4, UAS-GCaMP6f/UAS-IR62a; UAS-IR7c/+*
- *IR7c-Gal4/+; +/+; UAS-GCaMP7f/+*
- *IR7c-Gal4/+; +/UAS-IR62a; UAS-GCaMP7f/+*

Figure S1:

- *IR7c*^*Gal4*^*/IR7c*^*Gal4*^; *+/+; 20XUAS-CD8::GFP/+*
- *IR7c*^*Gal4*^*/IR7c*^*Gal4*^; *+/+; 20XUAS-CD8::GFP/20XUAS-CD8::GFP*

Figure S2:

- *IR7c*^*Gal4*^*/+; UAS-GCaMP7f/UAS-GCaMP7f; ΔIR62a/+*
- *IR7c*^*Gal4*^*/+; UAS-GCaMP7f/UAS-GCaMP7f; ΔIR62a/ΔIR62a*
- *IR7c*^*Gal4*^*/IR7c*^*Gal4*^; *UAS-GCaMP7f/+; ΔIR62a/ΔIR62a*

Figure S3:

- *IR7c*^*Gal4*^*/Gr66a-LexA::VP16; UAS-tdTomato/cyo* ; *LexAop-CD2::GFP/+*

Figure S4:

- *+/+; UAS-20xGCaMP6f/+; IR94e-Gal4, IR94e*^*LexA*^*/IR94e*^*LexA*^
- *+/+; UAS-20xGCaMP6f/+; UAS-IR7c, IR94e*^*LexA*^*/IR94e-Gal4, IR94e*^*LexA*^

### Tastants

The following tastants were used: sucrose, NaCl, MgCl_2_, CaCl_2_, caffeine, denatonium, acetic acid, serine, alanine, phenylalanine, glycine, NaBr, CsCl. All tastants were kept as 1M stocks and diluted as necessary for experiments.

### Immunohistochemistry

Immunofluorescence on labella was carried out as previously described^4^. For single labelling experiments, the primary antibody used was rabbit anti-GFP (1:1000, Invitrogen) and goat anti-rabbit Alexa 488 (1: 200, Invitrogen). For co-labeling experiments, chicken anti-GFP (1:1000, Abcam) and rabbit anti-RFP (1:200, Rockland Immunochemicals) were used as primary antibodies and goat anti-chicken Alexa 488 (1:200, Abcam) and goat anti-rabbit Alexa 647 (1:200, Thermo Fisher Scientific) were used as the secondary antibodies.

Brain immunofluorescence was performed as previously described^4^. Primary antibodies used were rabbit anti-GFP (1:1000, Invitrogen) and mouse anti-brp (1:50, DSHB #nc82). Secondary antibodies used were goat anti-rabbit Alexa 488 and goat anti-mouse Alexa 546 (1:200, Invitrogen).

All images were acquired using a Leica SP5 II Confocal microscope with a 25x water immersion objective. Images were processed in ImageJ^36^. For labellar analysis, confocal z-stacks of 2-8 labella were examined to identify neurons in each sensilla that were positive for the different drivers and where there was overlap.

### Behavioral assays

STROBE experiments were performed as previously described^24^. Mated females flies 2-3 days post eclosion were placed into vials containing 1 mL standard cornmeal food supplemented with 1 mM all-trans-retinal (Sigma-Aldrich) or an ethanol vehicle control for 2 days in the dark. Flies were starved for 24 hours on 1% agar supplemented with 1 mM all-trans-retinal, prior to experimentation. Both channels of the STROBE arenas were loaded with 4 μl of 100 mM sucrose. The STROBE software was started and single flies were placed into each arena via mouth aspiration. Experiments ran for 60 minutes, and the preference index for each fly was calculated as: (sips from Food 1 – sips from Food 2)/(sips from Food 1 + sips from Food 2).

Binary choice feeding assays were conducted as previously described^4^. Groups of 10 flies were starved on 1% agar for 24 hours prior to testing. For salt state experiments, flies aged 2– 5 days were sorted into groups of 10 and placed in salt fed (1% agar, 5% sucrose, and 10 mM NaCl) or salt deprived (1% agar and 5% sucrose) conditions for 3 days. Flies were 24-hr starved on 1% agar for the salt deprived group or 1% agar and 10 mM NaCl for the salt fed group. For all binary choice preference tests, flies were shifted into testing vials containing six 10 μL drops that alternated in color. Each drop contained the indicated concentration of tastant in 1% agar with either blue (0.125mg/mL Erioglaucine, FD and C Blue#1) or red (0.5mg/mL Amaranth, FD and C Red#2) dye. Each time a binary choice experiment was run, approximately half the replicates were done with the dye swapped to control for any dye preference. Flies were allowed to feed for 2 hours in the dark at 29°C before being frozen at −20°C. A dissection microscope was used to score the color of the abdomen as red, blue, purple, or no color. Preference Index (PI) was calculated as ((# of flies labeled with tastant 1 color) - (# of flies labeled with tastant 2 color))/ (total # of flies with color) and accounted for any flies that were lost in vial transferal and those that did not eat. Any vials with <7 flies or <30% of flies feeding were excluded.

### Calcium Imaging

Calcium imaging experiments were performed as previously described^4^. Prior to *in vivo* GCaMP imaging of the GRN axon terminals, flies were briefly anesthetized with CO_2_ and placed in a custom chamber. Nail polish was used to secure the back of the neck and a little wax was applied to both sides of the proboscis in an extended position, covering the maxillary palps without touching the labellar sensilla. After 1 hr recovery in a humidity chamber, antennae were removed along with a small window of cuticle to expose the SEZ. Adult hemolymph-like (AHL) solution (108 mM NaCl, 5 mM KCl, 4 mM NaHCO3, 1 mM NaH2PO4, 5 mM HEPES, 15 mM ribose, 2mM Ca^2+^, 8.2mM Mg^2+^, pH 7.5) was immediately applied. Air sacs and fat were removed and the esophagus was clipped and removed for clear visualization of the SEZ.

A Leica SP5 II Confocal microscope was used to capture GCaMPf fluorescence with a 25x water immersion objective. The SEZ was imaged at a zoom of 4x, line speed of 8000 Hz, line accumulation of 2, and resolution of 512 × 512 pixels. Pinhole was opened to 2.86 AU. For each taste stimulation, 20 total seconds were recorded. This consisted of 10 s baseline, 5 s stimulation, 5 s post-stimulation. A pulled capillary filed down to fit over both labellar palps was filled with tastant and positioned close to the labellum with a micromanipulator. For the stimulation, the micromanipulator was manually moved over the labellum and then removed from the labellum after 5 s. The stimulator was washed with water in between tastants of differing solutions. Salts were applied in order of increasing concentration and all solutions were applied in random order to control for potential inhibitory effects between modalities.

The maximum change in fluorescence (peak ΔF/F_0_) for peaks was calculated using peak intensity (average of 3 time points) minus the average baseline intensity (10 time points), divided by the baseline. ImageJ was used to quantify fluorescence changes and create heatmaps.

### Statistical Analysis

Statistical analyses were performed with GraphPad Prism 6 software. The number of biological replicates using different flies for each experiment and the statistical test performed is given in the figure legend. Sample sizes were determined ahead of experimentation based on the variance and effect sizes observed in prior experiments of similar types. Experimental conditions and controls were run in parallel. Data from calcium imaging experiments were excluded if there was too much movement during the stimulation to reliably quantify the response or if there was no response to a known, robust, positive control. The data from individual flies was removed from STROBE analyses if the fly did not pass a set of minimum sip threshold (15), or the data showed signs of a technical malfunction.

## Key Resources Table

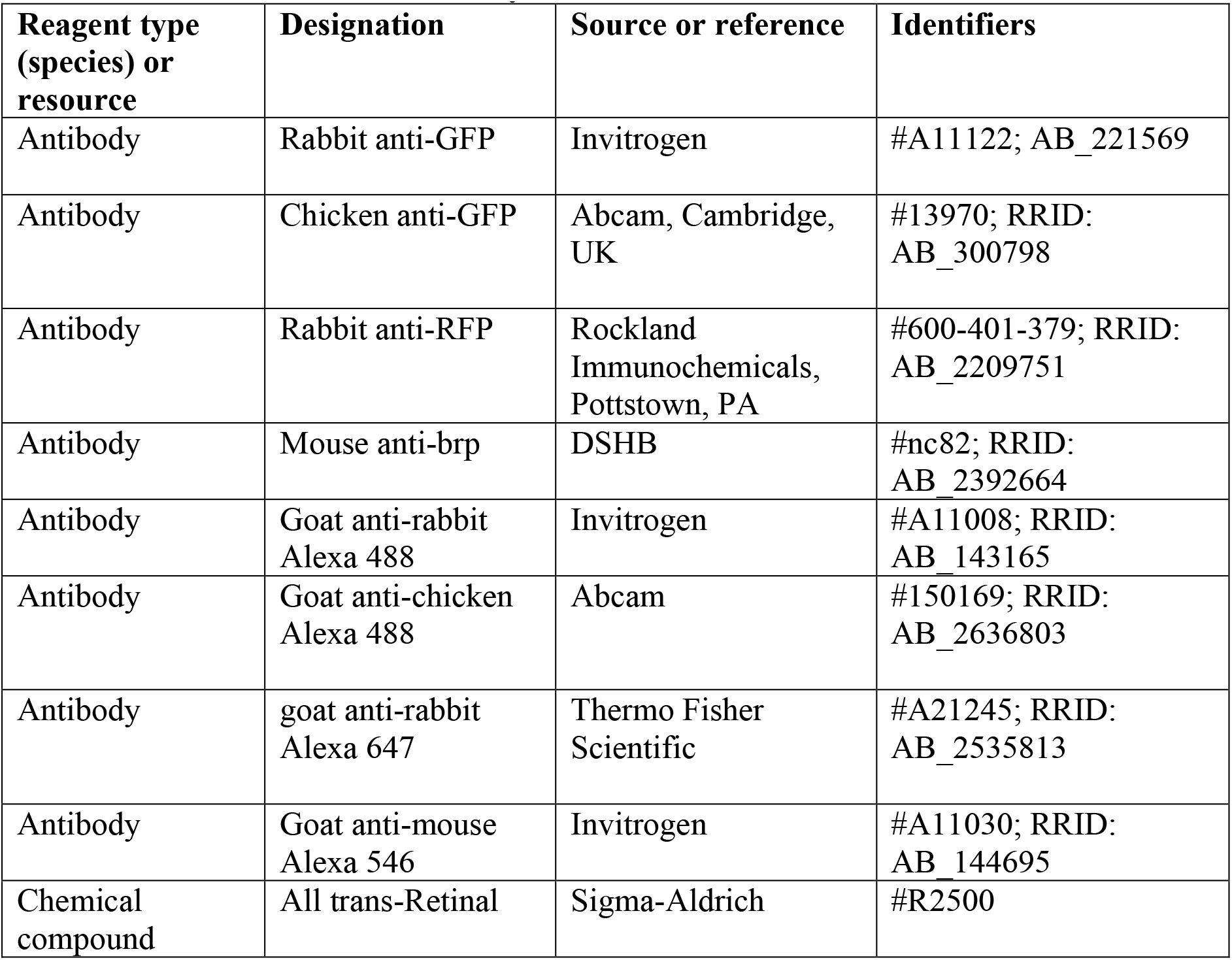
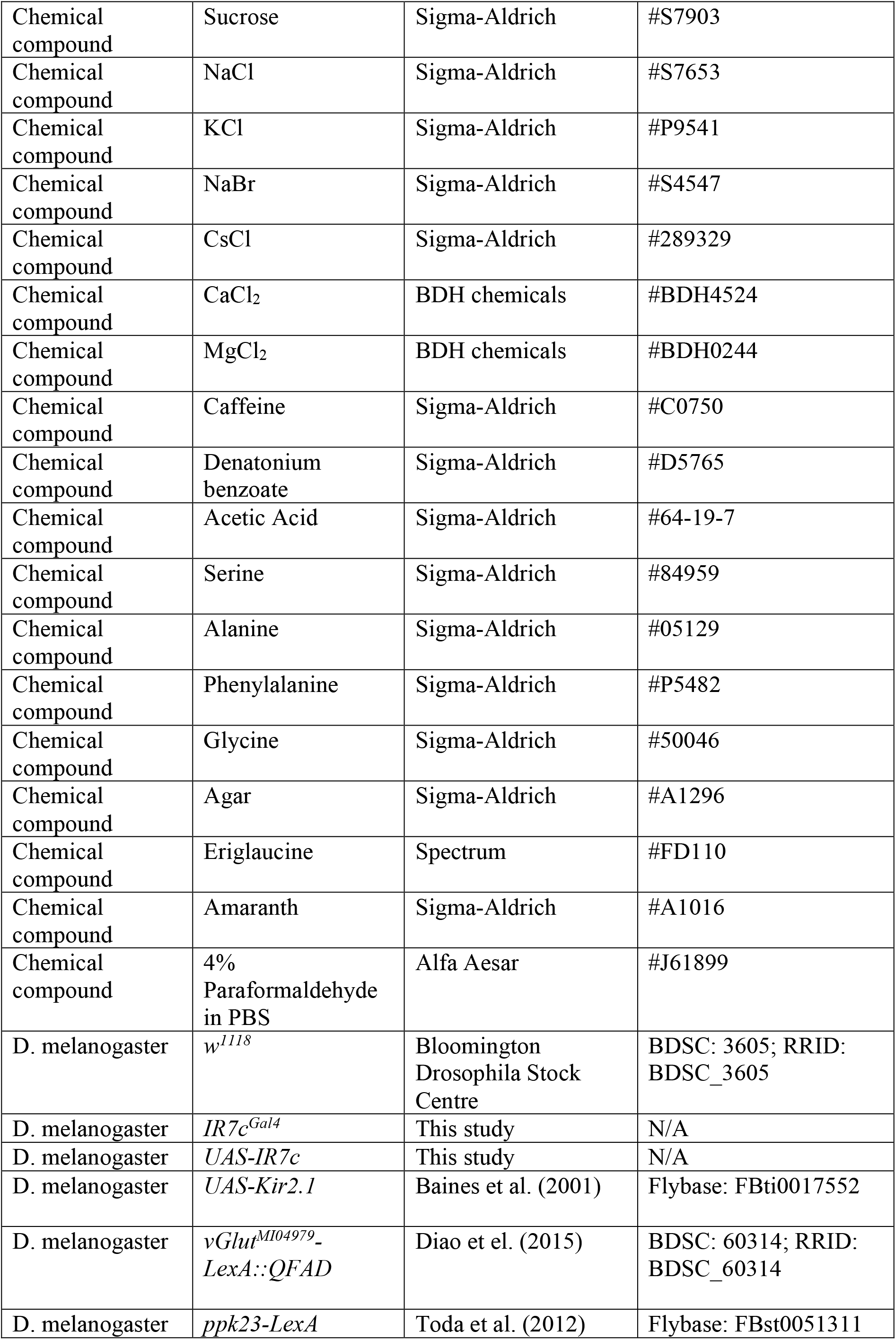
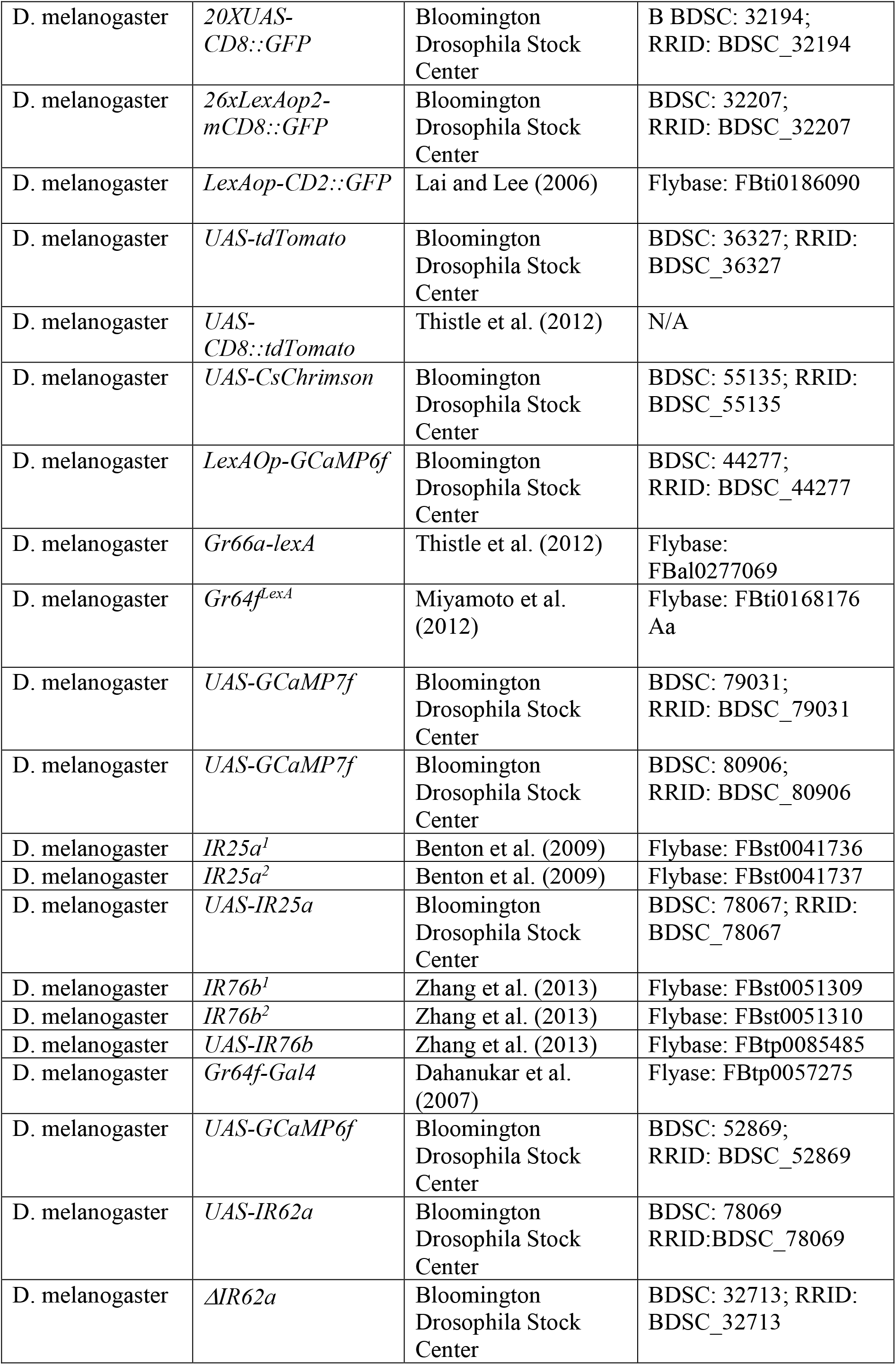
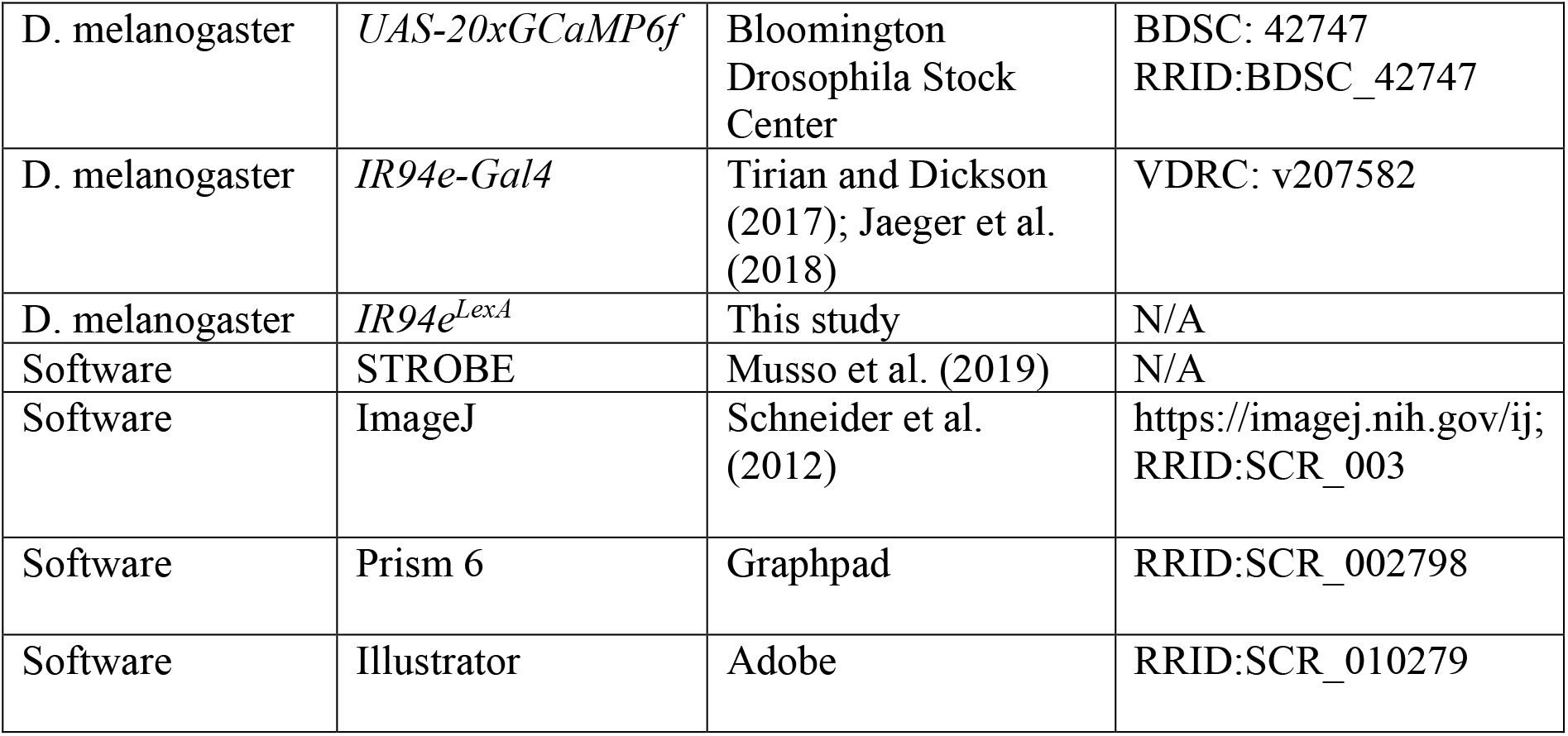

## SUPPLEMENTAL FIGURES

**Figure S1:**
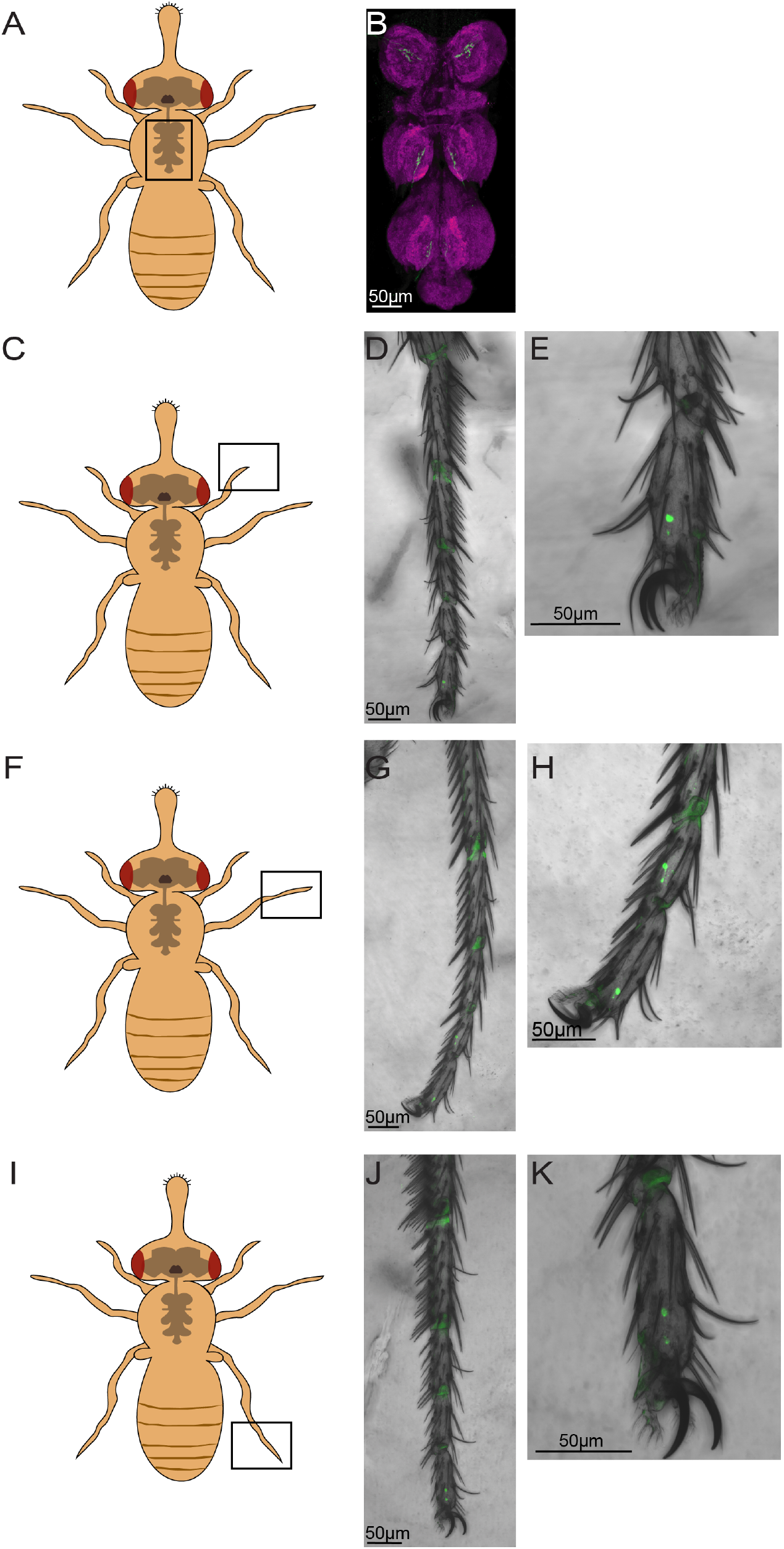
Characterization of IR7c GRNs in the fly VNC and tarsi. Related to Figure 1. **(A)** Schematic representation of VNC position. **(B)** Immunofluorescent detection of projections to the VNC labeled by *IR7c*^*Gal4*^ driving CD8::GFP. **(C-K)** Schematic representations of foreleg (C), midleg (F), and hindleg (I) position, with immunolabeling of CD8::GFP expressed in foreleg (D,E), midleg (G,H), and hindleg (J,K).

**Figure S2:**
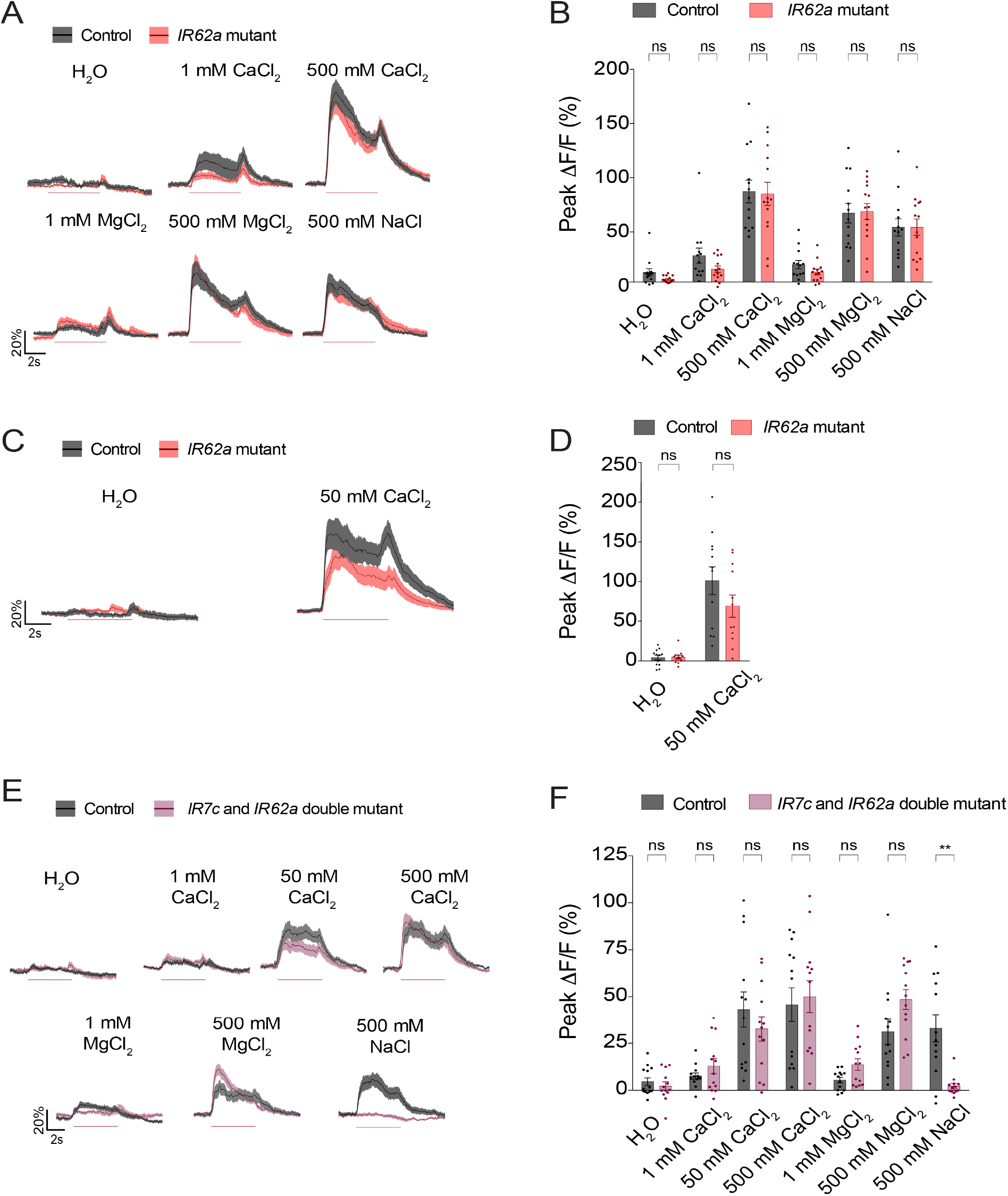
Divalent salt sensing in IR7c GRNs is mediated independently of both IR7c and IR62a. Related to Figure 3. **(A, C)** Traces of GCaMP7f signal in IR7c GRNs of *IR62a* mutants (red) and heterozygous controls (grey), following stimulation with the indicated tastants. **(B, D)** Peak fluorescence changes during each stimulation. n = 12-13 for each stimulus. **(E)** Traces of GCaMP7f signal in IR7c GRNs of *IR7c* and *IR62a* double mutants (pink) and isogenic controls (grey), following 5 s stimulation (red line) with the indicated tastants. **(F)** Peak fluorescence changes during each stimulation. n = 13 for each stimulus. All bars and traces represent mean ± SEM. Asterisks indicate significant difference between control and mutant responses for each stimulus by two-way ANOVA with Sidak’s post hoc test, **p<0.01.

**Figure S3:**
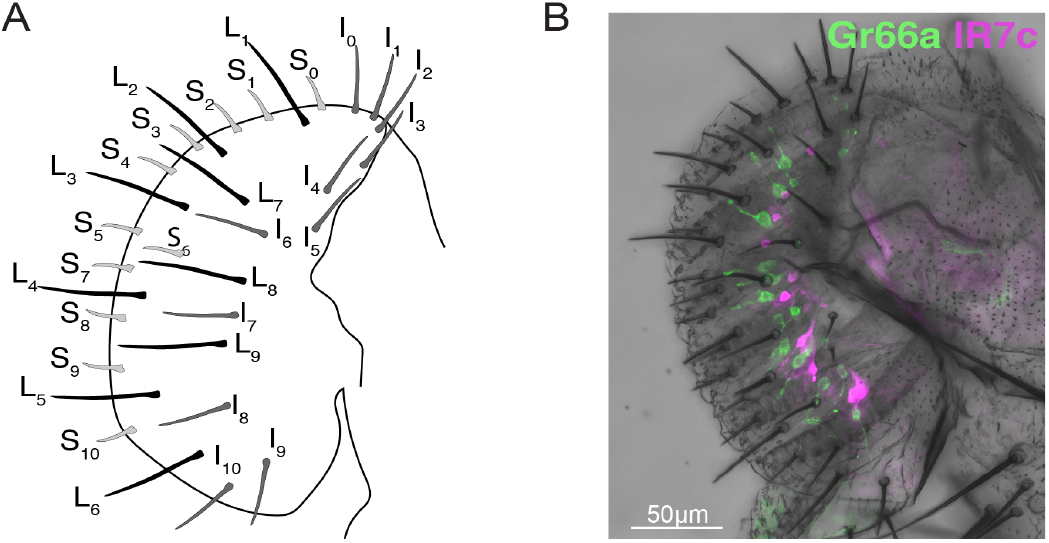
Co-labelling of IR7c GRNs with Gr66a GRNs. Related to Figure 4. **(A)** A schematic representation of the fly labellum and its sensillum identities. **(B)** *IR7c*^*Gal4*^ driving tdTomato (magenta) with GFP (green) under the control of *Gr66a-LexA*.

**Figure S4:**
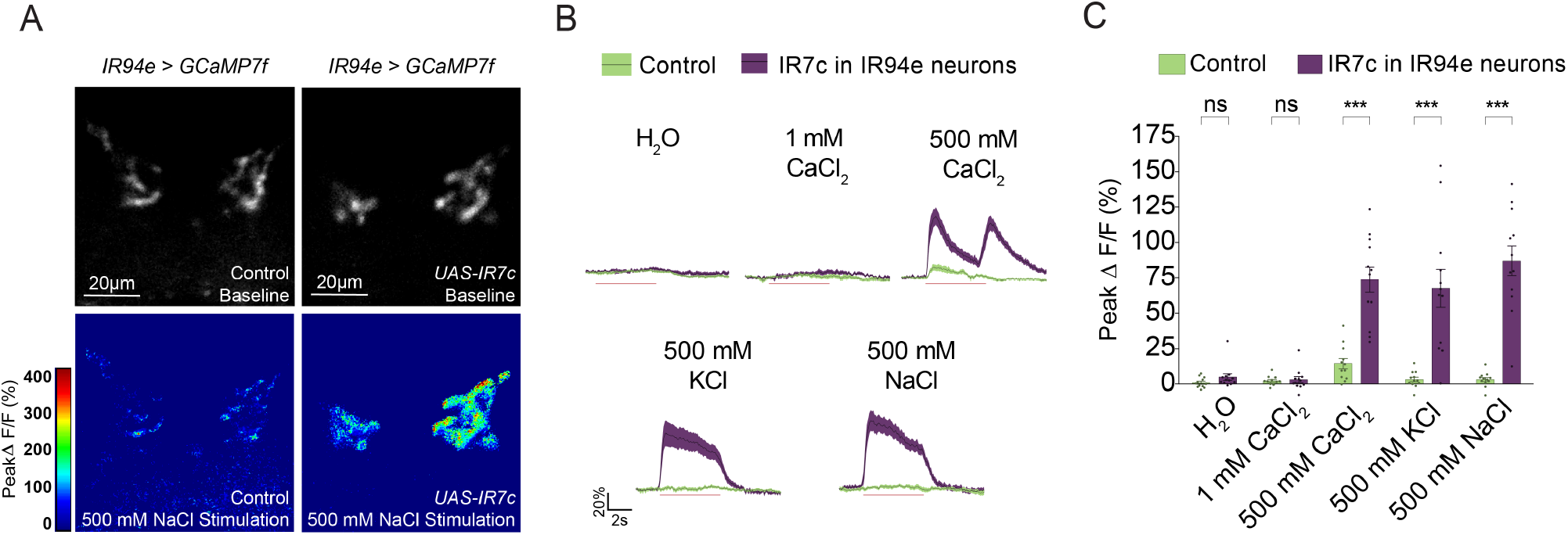
Ectopic IR7c expression in IR94e GRNs confers high salt sensitivity. Related to Figure 6. **(A)** Representative heatmaps of 500 mM NaCl-evoked activity in IR94e GRNs from an isogenic control (left) and a fly with IR7c expression under the control of *IR94e-Gal4* (right) both within an *IR94e* mutant background. **(B)** Traces of GCaMP6f signal in IR94e GRNs of flies with IR7c expression under the control of *IR94e-Gal4* (purple) and isogenic controls (green), following stimulation with the indicated tastants. Trace lines and shaded regions represent mean ± SEM. **(C)** Peak fluorescence changes during each stimulation. Bars represent mean ± SEM, n = 12 for each stimulus. Green dots (control), and purple dots (flies with IR7c expression under the control of *IR94e-Gal4*) indicate values for individual replicates. Asterisks indicate significant difference between responses of the two genotypes tested for each stimulus by two-way ANOVA with Sidak’s post hoc test, ***p<0.001.

## Notes

### Competing Interest Statement

The authors have declared no competing interest.

